# Ecology and environment predict spatially stratified risk of H5 highly pathogenic avian influenza clade 2.3.4.4b in wild birds across Europe

**DOI:** 10.1101/2024.07.17.603912

**Authors:** Sarah Hayes, Joe Hilton, Joaquin Mould-Quevedo, Christl A. Donnelly, Matthew Baylis, Liam Brierley

## Abstract

Highly pathogenic avian influenza (HPAI) represents a threat to animal and human health, with the ongoing H5N1 outbreak within the H5 2.3.4.4b clade being the largest on record. However, it remains unclear what factors have contributed to its intercontinental spread. We use Bayesian additive regression trees, a machine learning method designed for probabilistic modelling of complex nonlinear phenomena, to construct species distribution models (SDMs) for HPAI clade 2.3.4.4b presence. We identify factors driving geospatial patterns of infection and project risk distributions across Europe. Our models are time-stratified to capture both seasonal changes in risk and shifts in epidemiology associated with the succession of H5N6/H5N8 by H5N1 within the clade. While previous studies aimed to model HPAI presence from physical geography, we explicitly consider wild bird ecology by including estimates of bird species richness, abundance of specific taxa, and “abundance indices” describing total abundance of birds with high-risk behavioural traits. Our projections of HPAI clade 2.3.4.4b indicate a shift in persistent, year-round risk towards cold, low-lying regions of northwest Europe associated with H5N1. Methodologically, we demonstrate that while most variation in risk can be explained by climate and physical geography, adding host ecology is a valuable refinement to SDMs of HPAI.

## Introduction

Avian influenza is a highly contagious viral disease caused by avian influenza type A viruses. It can have devastating effects on animal health and livestock economics with losses reaching $billions in the USA alone ^1^, but also poses a threat to human public health and has pandemic potential. Spillover of avian influenza from birds to mammals is known to occur, but most spillover events are associated with little or no ongoing transmission. However, recent reports of mass mortality events in marine mammal populations in South America ^2–4^ and farmed mink in Spain ^5^ sparked concerns surrounding the potential for mammal-to-mammal transmission. The discovery of HPAI infection in cattle in North America in 2024 ^6,7^ and subsequent instances of onward transmission to farm workers ^8^ have prompted high-priority surveillance responses considering the threat posed by potential human adaptation of the virus.

Whilst highly pathogenic avian influenza (HPAI) viruses may infect both domestic and wild birds, they are defined as those that cause severe disease in young poultry chicks intravenously inoculated in a laboratory setting, or those that possess multiple basic amino acids in the haemagglutinin cleavage site, which has been associated with high virulence. HPAI viruses can arise from changes to low pathogenic avian influenza (LPAI) viruses during circulation within poultry and are often detected from the resulting severe disease. HPAI is typically restricted to subtypes containing the H5 or H7 haemagglutinin glycoprotein (HA). Since 2002, symptomatic HPAI in wild birds has been increasingly recognised ^9–13^. The HPAI virus strain A/goose/Guangdong/1/1996 (H5N1) (gs/Gd) first emerged in geese in southeast Asia in 1996 (14) and was reported to cause wild bird mortality several years later ^10^. By late 2005, gs/Gd HPAI viruses were recognised in Europe, with the migration of wild birds implicated in this long-distance transmission ^15,16^. Since then, genetic divergence and reassortment events have led to multiple European outbreaks of various clades within the gs/Gd HPAI lineage in both domestic and wild birds ^13,17^.

Since emerging out from the Qinghai Lake region, clade 2.3.4.4b has arisen as the dominant clade in Eurasia, with significant European outbreaks of H5N8 genotype viruses in 2016-17 and 2020-21 ^18–20^. A new reassortant H5N1 genotype then emerged in Europe around late 2020 ^19^, containing 2.3.4.4b HPAI HA and matrix protein (MP) genes, but LPAI-origin neuraminidase and internal genes ^18^. This reassortant has since significantly expanded both in geography ^17,21^ and host range ^19^, affecting a much wider diversity of wild species, including gannets (*Sulidae*), pelicans (*Pelicanidae*), terns (*Sterninae*), and skuas (*Stercorariidae*).

Phylodynamic analyses also show wild birds experience longer duration of circulation, wider spatial dissemination, and greater likelihood of seeding infections of 2.3.4.4b viruses compared to domestic poultry ^22^ or wild birds infected with previous H5Nx clades ^18,19^, widening the possible palette of future reassortment risks in the wild. The apparent broader host susceptibility and persistence of H5N1 2.3.4.4b in particular is thought to be driven by further reassortment events with endemic gull H13 LPAI viruses ^20^. Therefore, there is a critical need to better understand the changing role of wild birds in the transmission and maintenance of avian influenza viruses.

There has been much interest in which species contribute to population maintenance of influenza viruses. In 2006, six HPAI H5N1 viruses were isolated from apparently healthy migratory ducks in China ^23^ and recent infection studies have suggested that mallard ducks (*Anas platyrhynchos)* may shed H5N1 clade 2.3.4.4b virus whilst asymptomatic ^24,25^. In addition to risks associated with particular host species, wider functional ecology of hosts has been cited as an important but neglected factor in the epidemic transmission of viral diseases of wild birds ^26,27^. Behaviours that increase bird density and host-to-host direct and indirect contact rates such as colony breeding and pre-migratory congregation may be associated with increased risk of exposure ^28^. Migration also places a high physiological demand on birds, which may disrupt immunity ^29^ and thus increase susceptibility to infection, although evidence for this is somewhat conflicting ^28^. For LPAI, viruses in water can remain infectious for more than 30 days at 0°C with spread predominantly via the faecal-oral route therefore dependent on foraging behaviours.

The high prevalence of LPAI viruses in dabbling ducks may be in part due to exposure via surface water feeding ^30,31^. However, HPAI viral shedding has also been linked to the respiratory route ^32–35^ which may alter dependency of transmission on foraging behaviours. Even further unknown is the specific ecology of HPAI clade 2.3.4.4b viruses and whether it may explain their apparent diversity of susceptible hosts.

Species distribution models (SDMs) make predictions across geographic space and time using a set of predictor variables. They have traditionally been used in ecology to predict the distribution of plant and animal species using environmental variables, but more recently have been applied to infectious disease^36–40^. SDMs have been used for modelling avian influenza viruses in domestic birds ^41–47^, wild birds ^48–54^, and both combined ^55,56^. Models have been produced at local ^52–54^ and continental resolutions ^48–51^ but few have adopted more recent approaches to integrate host ecology at large spatial scales ^26^, instead using only environmental predictors. Notable exceptions include Belkhiria *et al.* ^52^, who included Important Bird and Biodiversity Areas as model predictors to act as a proxy for the presence of migratory birds, whilst Moriguchi *et al.* ^53^ included population sizes of migratory and dabbling ducks. Hill *et al.* ^57^ used poultry holding densities, poultry species-specific risk estimates, and wild bird population sizes to inform a spatial mechanistic model of incursion into poultry farms in the UK, but did not include physical geography in their model. Huang *et al.* ^50^ explored more sophisticated ecological measures of host community composition on HPAI H5N1 across Europe, incorporating features such as species richness and abundance of waterbirds in general and of specific waterbird high-risk hosts, as well as phylogenetic relatedness to those high-risk hosts. However, none of these previous studies combine a continental scale of analysis, ecological traits of the host, and a direct comparison between the current outbreak of H5N1 HPAI with that of previously circulating H5 HPAI lineages within clade 2.3.4.4b.

The aim of this study is to map and understand the geospatial risk of HPAI presence across Europe using an SDM approach. Our modelling builds on previous studies which focus on physical geography by explicitly incorporating ecological covariates including avian species richness, abundance of avian species with specific behavioural risk factors, and abundance of specific high-risk avian taxa. We train a Bayesian additive regression trees (BART) model on H5 HPAI detections of clade 2.3.4.4b since 2016 and use data on the physical geography and ecology of over 600 wild bird species of Europe to estimate the probability of HPAI clade 2.3.4.4b presence at a 10km resolution. Although we primarily use models to identify specific locations at high risk of acting as epicentres for future HPAI outbreaks, we also calculate variable importance and partial dependence to interrogate which specific risk factors are highly predictive of different subtypes of H5 clade 2.3.4.4b and why, including novel contributions from host ecological data. We sense-check our models by comparing their risk projections with data on domestic bird HPAI detections, as introduction of HPAI into domestic poultry may result from contact with infected wild birds or wild bird-derived fomites, and so we expect domestic poultry cases to be more likely to occur in areas where we predict high risk of HPAI in wild birds.

## Methods

### Data assembly and season definition

We assembled a set of geospatial covariate rasters at a 10km spatial resolution. This resolution was chosen to smooth potential error in geolocated coordinates of positive HPAI records given the wide-ranging movement of wild bird hosts, and for close alignment with the native resolution of many covariates (Table 1). We used the EPSG:3035 coordinate reference system, which is intended for spatial statistical analysis across continental Europe ^58^. To account for seasonal changes in risk, we assembled data for four seasonal periods as an overall calendar proxy for European bird behavioural cycles: non-breeding season, 30th November - 28th (or 29th) February (calendar days 334 - 59); pre-breeding migration, 1st March - 6th June (days 60 - 157); breeding season, 7th June - 9th August (days 158 - 221); and post-breeding migration, 10th August - 29th November (days 222 - 333).

**Table 1.** Environmental covariates included in the models.

| Topographic Covariates | Details | Resolution of data source | Year of data | Source |
| --- | --- | --- | --- | --- |
| Elevation | Minimum elevation (metres above sea level) | 30 arc seconds | GTOP030 - 1999<br>CGIAR-SRTM - 2018 | Geodata package R https://cran.r-project.org/web/packages/geodata/geodata.pdf |
| Elevation | Maximum elevation (metres above sea level) | 30 arc seconds | GTOP030 - 1999<br>CGIAR-SRTM - 2018 | Geodata package R https://cran.r-project.org/web/packages/geodata/geodata.pdf |
| Elevation | Difference between minimum and maximum elevation (m) | 30 arc seconds | GTOP030 - 1999<br>CGIAR-SRTM - 2018 | Processed using minimum and maximum elevation listed above |
| Elevation | Modal elevation (metres above sea level) | 30 arc seconds | GTOP030 - 1999<br>CGIAR-SRTM - 2018 | Geodata package R https://cran.r-project.org/web/packages/geodata/geodata.pdf |
| Normalised Difference Vegetation Index (NDVI) | Mean over the season. Normalised to limit values to between -1 and 1 | 1km | 2022 | Modis https://modis.gsfc.nasa.gov/data/dataproduct/mod13.php |
| Land cover | One raster layer for each of the 17 land cover categories | 0.5km | 2022 | Modis https://lpdaac.usgs.gov/products/mcd12q1v061/ |
| Distance to coast | Distance from centre of raster cell to nearest coast (km) |  | N/A | Calculated directly from map data |
| Distance to inland water | Distance from centre of raster cell to nearest permanent | 30 arc seconds | 2004 | Calculated using data from Global |
|  | inland water (m) |  |  | Lakes and Wetland Database<br>https://www.worldwildlife.org/publications/global-lakes-and-wetlands-database-lakes-and-wetlands-grid-level-3 |
| **Bioclimatic Covariates** |  |  |  |  |
| Relative humidity | Seasonal mean of daily midday (local time) relative humidity measured at 2 metres (%) | 0.1 degrees | 2022 | Copernicus<br>https://cds.climate.copernicus.eu/cdsapp#!/dataset/sis-agrometeorological-indicators?tab=overview |
| Temperature | Seasonal weighted mean of the month-wise difference in the minimum temperature and maximum temperature (degrees Celsius) | 2.5 arc minutes | 2018 | WorldClim<br>https://www.worldclim.org/data/monthlywth.html |
| Temperature | Seasonal weighted mean of monthly mean temperatures (degrees Celsius)<br>(Mean monthly temperature for each month calculated using: Mean temperature = Minimum temperature + diurnal range/2) | 2.5 arc minutes | 2018 | WorldClim<br>https://www.worldclim.org/data/monthlywth.html |
| Temperature | Seasonal temperature variation (degrees Celsius)<br>(Difference between the maximum and minimum of mean monthly temperature values across months majority-represented within the season) | 2.5 arc minutes | 2018 | WorldClim<br>https://www.worldclim.org/data/monthlywth.html |
| Precipitation | Seasonal weighted mean of the monthly total precipitation (mm) | 2.5 arc minutes | 2018 | WorldClim<br>https://www.worldclim.org/data/ |
|  |  |  |  | monthlywth.html |
| Zero-degree isotherm | Seasonal mean of daily zero-degree isotherm (metres above sea level) | 0.25 degrees | 2022 | Copernicus<br>https://cds.climate.copernicus.eu/cdsapp#!/dataset/reanalysis-era5-single-levels?tab=overview |
| Zero-degree isotherm | Number of days the zero-degree isotherm was below 1 metre at midday at Coordinated Universal Time (UTC) | 0.25 degrees | 2022 | Copernicus<br>https://cds.climate.copernicus.eu/cdsapp#!/dataset/reanalysis-era5-single-levels?tab=overview |
| **Livestock demography** |  |  |  |  |
| Chicken density | Headcount of animals | 5 arc minutes | 2010 | Gridded livestock of the world 3<br>https://www.fao.org/livestock-systems/global-distributions/chickens/en/ |
| Duck density | Headcount of animals | 5 arc minutes | 2010 | Gridded livestock of the world 3<br>https://www.fao.org/livestock-systems/global-distributions/ducks/en/ |

These periods were chosen by querying seasonal status for each bird species listed as present in Europe within the 2022 data release of eBird Status and Trends ^59^. These data describe numerical calendar weeks of each individual bird species’ life cycle season (“breeding”, “nonbreeding”, “post-breeding migration”, and “pre-breeding migration” seasons, or “resident” for non-migratory species). We defined aggregated proxy behavioural seasons such that a plurality of migratory bird species are in the given season at the given time (Supplemental Methods S1, Supplemental Figure S1A). For example, from 30th November - 28th February, the majority seasonal status among migratory birds is ‘non-breeding’. At the species level, our behavioural seasons are intended to capture the modal behaviour of European seasonal birds.

### Environmental covariates

We selected a set of environmental covariates consistent with those reported to be important in avian influenza dynamics ^48–54,60^. Covariates were categorised as topographical, bioclimatic or livestock demography and are listed in Table 1. There was a large amount of variability in the resolution and coordinate reference systems of the different data sets used. Where calculations were required these were done at the original resolution prior to reprojection. Rasters were then reprojected to the EPSG:3035 coordinate reference system and the resolution converted to 10km using bilinear interpolation for continuous covariates and nearest neighbour for categorical covariates.

#### Topographical

Elevation data were sourced using the R package “geodata” v0.5-8 ^61^ which combines Shuttle Radar Topography Mission (SRTM) data and Global 30 Arc-Second Elevation (GTOPO30) data. Elevation data were processed to extract the minimum, maximum and modal elevation for each of the 10km resolution cells of our project area. A fourth elevation raster containing the difference between the minimum and maximum values recorded within each of the cells was also produced.

The normalised difference vegetation index (NDVI) is a measure of the vegetation canopy greenness and acts as a marker of vegetation density. NDVI data were sourced from NASA’s Moderate Resolution Imaging Spectroradiometer (MODIS) data in 16-day periods at a 1km resolution. The data were grouped to reflect the seasons as closely as possible by assigning each 16-day period to the season the majority of its days fell within. A mean NDVI for each aggregated bird behavioural season was calculated and standardised to limit the range between -1 and 1.

Data on land cover were sourced from MODIS Land Cover Type (MCD12Q1) Version 6.1. Land cover was categorised as one of 17 different land cover types numbered 1-17. During reprojection, the nearest neighbour method was used to create a raster at 10km resolution. Separate rasters for each of the land cover types are then created with the value of the cell representing whether or not that land cover type is assigned to that cell.

Inland water bodies were identified using the Global Lakes and Wetlands database provided by the World Wildlife Fund. We excluded intermittent (i.e., non-permanent) lakes and wetlands with the resulting raster layer containing distance to any permanent water body of any type.

#### Bioclimatic

Temperature and precipitation data were obtained from WorldClim historical monthly data for the most recent year available at initial download (2018) at a 2.5 arc minute resolution. Temperature data are entered as seasonal weighted mean of monthly mean temperatures (weighting by number of days of each month represented in the behavioural season), short-term variability in temperature taken as weighted mean of the month-wise differences in minimum and maximum temperatures, and the overall variation in mean temperature over the season (calculated by taking the difference between the minimum and the maximum of the mean temperatures across all months with at least half their days represented in the respective season). Precipitation is entered as the mean (weighted by days falling in season, as for temperature) of the monthly values for total precipitation for each of the months of the season.

Relative humidity data were downloaded from the European Centre for Medium Range Weather Forecasts (ECMWF) Copernicus dataset, obtained by the ERA-Interim reanalysis of monthly observations from 1979 to the present. Data reflect the daily relative humidity at 2 metres above ground level at midday local time. Mean seasonal humidity was calculated as the mean of the daily humidity values within the season.

Daily zero-degree isotherm data were also obtained from Copernicus and reflect the height above the Earth’s surface where the temperature in Celsius passes from positive to negative values at the specified time. Seasonal mean zero-degree isotherm and the number of days within the season where the zero-degree isotherm was below 1 metre at midday (UTC) are included in the model.

#### Livestock demography

Data regarding the density of chickens and ducks were obtained from Gridded Livestock of the World version 3. Version 3 was used for both species due to large amounts of missing data for ducks in version 4.

### Ecological covariates

We constructed raster layers of ecological risk factors by combining species-level risk factors, estimates of global species population and relative spatial abundance data into a single index of ‘species-trait abundance’. To reflect the key role of migratory behaviour in wild bird HPAI epidemiology ^22,27,62,63^ we obtained binary species-level indicators specifying whether a given species was migratory (defined as having ‘Full Migrant’ classification) and/or congregative from the IUCN Red List ^64^. To capture bird diet and foraging behaviour, we took estimates from EltonTraits ^65^ of the percentage contribution of specific food sources and foraging behaviours to each species’ total diet and time spent foraging. Of the food sources considered in EltonTraits, we considered plants and endothermic vertebrates (mammals and birds). Of the foraging behaviours, we considered feeding within 2m of water surfaces, feeding below the water surface at depths exceeding 2m, and scavenging. Evolutionary relatedness between hosts is a well-established predictor of infection potential in various systems ^66,67^, including avian influenza ^68^. To capture this effect of phylogenetic structure in susceptibility across avian host species, we downloaded 100 posterior estimates of the bird phylogenetic tree from BirdTree ^69^ and used the function cophenetic.phylo from the R package ape, v5.8 ^70^ to calculate a matrix of pairwise phylogenetic distances between bird species. We used the CLOVER database ^71^ to identify known avian influenza host species, and for each species in the BirdTree phylogeny calculated the minimum phylogenetic distance to a known host species. Based on preliminary analysis we converted this to an indicator variable specifying for each species whether the phylogenetic distance to any known avian influenza host species met a threshold value of 50 million years since lineage divergence. Taxonomic conventions differed between data sources, with disagreements over both species names and distinctions between species and subspecies. To allow for cross-referencing between data sources with differing taxonomic conventions we performed a semi-automated rationalisation process using the AVONET database of bird species names ^72^, detailed in Supplemental Methods S1. A complete list of ecological covariates used in our models is given in Table 2.

**Table 2.** Ecological covariates included in the models.

| Covariate | Total abundance or behaviour-weighted? | Species-level source |
| --- | --- | --- |
| *Anatinae* (“dabbling ducks”) | Total | eBird<br>https://science.ebird.org/en/status-and-trends |
| *Anserinae* (swans and geese) | Total | eBird<br>https://science.ebird.org/en/status-and-trends |
| *Ardeidae* (herons) | Total | eBird<br>https://science.ebird.org/en/status-and-trends |
| *Arenaria/Calidris* (turnstones and sandpipers) | Total | eBird<br>https://science.ebird.org/en/status-and-trends |
| *Aythya* (“diving ducks”, or pochards) | Total | eBird<br>https://science.ebird.org/en/status-and-trends |
| *Laridae* (gulls) | Total | eBird<br>https://science.ebird.org/en/status-and-trends |
| Percentage time spent feeding within 2m of water surface | Weighted | EltonTraits<br>https://doi.org/10.6084/m9.figshare.c.3306933.v1 |
| Percentage time spent feeding >2m below water surface | Weighted | EltonTraits<br>https://doi.org/10.6084/m9.figshare.c.3306933.v1 |
| Percentage diet plants | Weighted | EltonTraits<br>https://doi.org/10.6084/m9.figshare.c.3306933.v1 |
| Percentage diet scavenging | Weighted | EltonTraits<br>https://doi.org/10.6084/m9.figshare.c.3306933.v1 |
| Percentage diet endothermic vertebrates | Weighted | EltonTraits<br>https://doi.org/10.6084/m9.figshare.c.3306933.v1 |
| Congregative | Total | IUCN Red List |
|  |  | https://www.iucnredlist.org/ |
| Migratory | Total | IUCN Red List<br>https://www.iucnredlist.org/ |
| Below threshold phylogenetic distance to known host species | Total | BirdTree<br>https://birdtree.org/<br>CLOVER<br>https://github.com/viralemergence/clover |
| Species richness | N/A | eBird<br>https://science.ebird.org/en/status-and-trends |

We obtained estimates of species abundance in the form of percentage population at a 9km grid resolution from the spatiotemporal component of eBird Status and Trends. These estimates are calculated using species distribution models trained on checklist-based bird observation data submitted to the eBird citizen science platform, and, for a given species, take the form of 52 weekly geospatial raster layers specifying the percentage of the global population of a given species in each grid cell in each week of the year. The species modelled by eBird include only a subset of those known to be present in Europe; a total of 708 species as opposed to 944 species listed as being observed in Europe in Avibase ^73^. For each species in each behavioural season, eBird provides quality ratings on a scale of 0 (lowest quality) to 3 (highest quality).

We used the species-specific quality metrics and temporal boundaries to generate weekly quality measures, and for each aggregated behavioural season included only those species which had a quality rating ≥2 for the majority of the season. We defined the estimated abundance of a given species in a given grid cell for a given aggregated behavioural season to be the mean of the weekly abundance across all weeks in that season with quality rating ≥2. To convert from percentage population to absolute abundance we subsequently scaled each species’ abundance raster by the corresponding estimated global population size provided by Callaghan *et al.* ^74^, giving us an estimated absolute abundance raster. After discarding species with problematic taxonomic statuses and year-round low quality levels we were left with 617 of the 708 European species modelled by eBird Status and Trends.

We used these estimated absolute abundance layers to develop spatial raster layers representing indices of population-level ecological traits for use as covariates. Species richness was estimated by counting the total number of species in each grid cell that had a total estimated abundance greater than 1. For migratory behaviour, congregative behaviour, and phylogenetic host distance threshold, we created indices by adding together all the abundance layers for species which had a positive indicator for these factors. For the dietary and foraging behaviour risk factors, we summed all of the abundance layers weighted by the percentage of dietary and foraging behaviour allocated to the specific behaviour for each species in EltonTraits, to give a measure of relative population behaviour in each grid cell. We constructed covariate layers representing six specific bird taxa by adding together all the abundance layers for species in each. These taxa were families Anatinae, Anserinae, Ardeidae, Aythyini, Laridae, as well as genera *Arenaria* and *Calidris*, which were combined in a single layer due to their close relatedness and the small number of species in *Calidris* ^69^. These taxa were chosen based on high numbers of appearances in the CLOVER database ^71^ or previous identification as high-risk avian influenza species ^75^.

### Outcome data

Data on confirmed avian influenza cases in wild birds were obtained from publicly available data sources: The Food and Agricultural Organization of the United Nations’ (FAO) EMPRES-i and the World Animal Health Information System (WAHIS) database provided by the World Organization for Animal Health (WOAH). Data spanned from August 2016 until February 2024 and were filtered to H5 HPAI within the specified study area. Duplicate entries based on date and geographical coordinates were removed. Data entries that had the same date and geographical coordinates but were listed as different subtypes were retained within the data. Data were mapped to a 10km grid considering cells positive if they contained ≥1 HPAI record.

Based on patterns of epidemic peaks and subtype emergence in Europe (Figure 1), independent models were constructed to capture A) discrete H5 HPAI outbreaks within clade 2.3.4.4b of H5N8 and H5N6 from 2016 to 2021 and B) the extended H5N1 clade 2.3.4.4b epizootic, ongoing since 2021. Based on behavioural season boundaries, each set was divided into a training period, and a subsequent test period held out from training to assess model performance in near-future projections. For A) models were trained on cases from 10/8/2016 to 9/8/2020, capturing H5N8 and H5N6 events (Figure 1), before testing on a separate H5N8 outbreak from 10/8/2020 to 9/8/2021. For B) models were trained on H5N1 cases from 10/8/2021 to 28/2/2023 before testing on H5N1 cases from 1/3/2023 to 29/2/2024.

**Figure 1.**
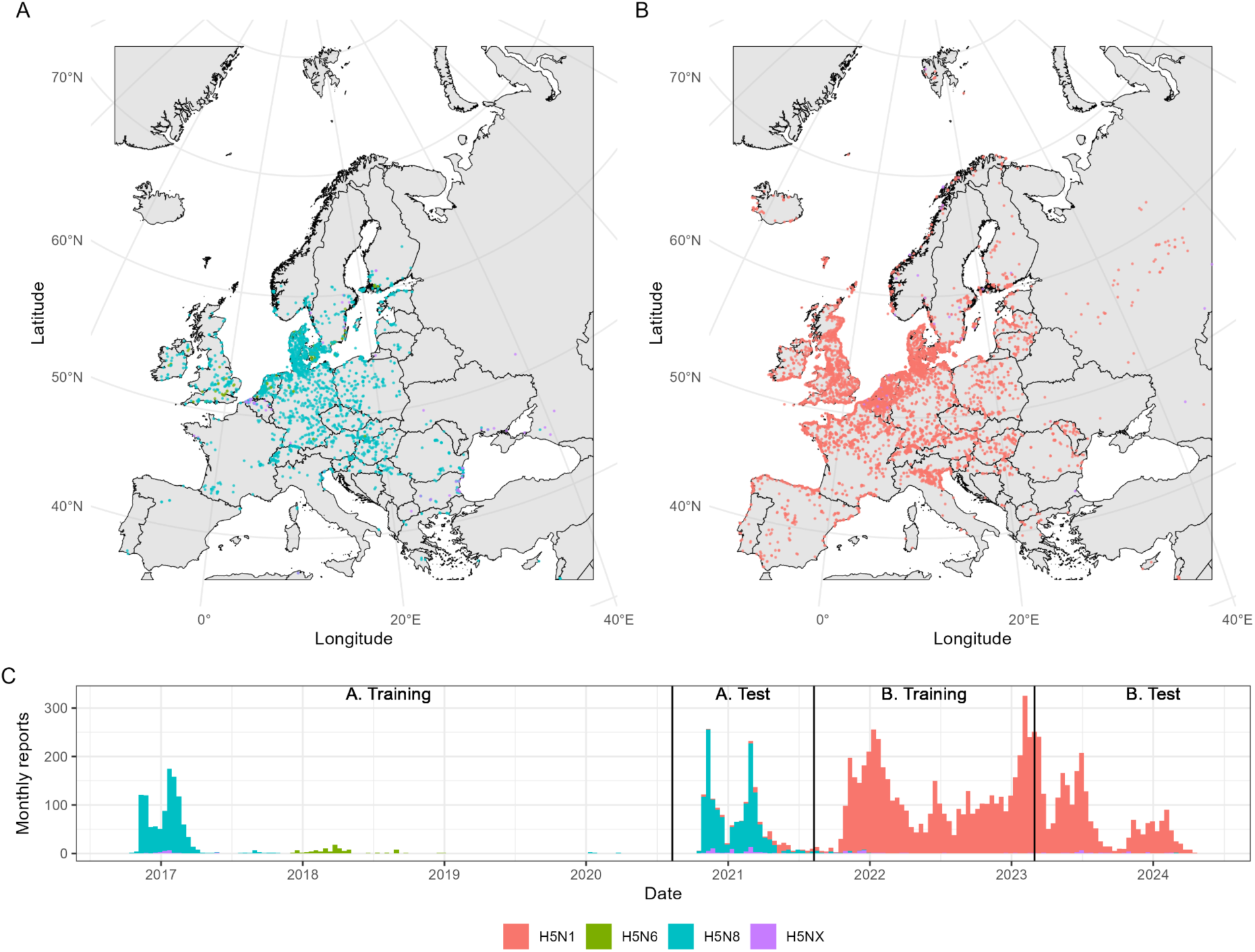
Wild bird H5 HPAI clade 2.3.4.4b incident data in Europe. Spatiotemporal distribution of incidents of H5 highly pathogenic avian influenza clade 2.3.4.4b reported in wild birds, harmonised from EMPRES-i and WAHIS databases. A) and B) Geolocated incident reports over time periods A (10/8/2016 - 9/8/2021) and B (10/8/2021 - 29/2/2024), respectively. C) Total independent incident reports per month with separate model training and test periods annotated. Colour denotes subtype recorded (H5N1; H5N6; H5N8; or other H5 where full subtype not explicitly labelled, denoted H5NX).

Training/test divisions were chosen to cover a minimum of one full bird behavioural season of each type (Supplemental Figure S1B).

### Pseudo-absence generation

SDM analyses are often challenged by lack of available absence data (i.e., true negatives) to train on alongside presences; for HPAI, few authorities systematically report their negative test results. Several approaches exist to handle this problem, including presence-only or ‘one-class’ classification and niche boundary estimation ^76,77^. We use an established solution in generating ‘pseudo-absence’ or background points, that can fulfil the role of negative training data by representing areas the modelled organism or disease could feasibly be present in but has never been observed in. It is strongly emphasised that pseudo-absences should be drawn to reflect the same biological restrictions and spatial sampling heterogeneities as presences ^76,78^. We expect influenza surveillance in wild birds to be restricted by both human population to report potential cases and physical accessibility to conduct sampling. Therefore, to approximate these processes, we obtained general citizen surveillance records of wild bird sightings.

We obtained all bird observations made by the public in eBird by downloading the eBird Basic Dataset (EBD) ^79^ before filtering to records within Europe using R package ′auk′, v0.7.0 ^80^. We then calculated total sightings (i.e., unique data-user-geolocation records, regardless of species sighted) per 10km grid cell over the study period 10/8/2016 - 29/2/2024 to create a spatial raster layer approximating bird sampling accessibility (Supplemental Figure S2). Pseudo-absences were then sampled from all cells that remained HPAI-free during the entire study period and were ≥ 25km from the nearest positive cell.

Pseudo-absences were drawn at an initial 1:1 ratio with positives for each season using function add_pseudoabsence() from R package ′ibis.iSDM′ v0.1.2 ^81^, weighting by normalised total eBird sightings. We therefore preferentially select cells with high human density and sampling accessibility yet have never reported HPAI as those we have stronger confidence in to represent true negatives. Pseudo-absences were generated jointly for training/test sets to avoid positive cells from one being considered pseudo-absences in the other.

### Data resampling

As a result of sampling patterns, spatial occurrence records for organisms or diseases are commonly concentrated in clusters, which can bias SDMs and their predictions. As in other computational fields, these biases can be mitigated by resampling training data with spatial data thinning being a popular approach ^82^. However at continental scales, this may retain records that are geographically diverse, but capture little environmental variation.

Therefore, we reduce training data redundancy by environmental thinning, which typically outperforms spatial thinning ^83,84^. Considering environmental traits as an n-dimensional space and dividing each trait into a given number of bins, simple random stratified sampling can be conducted based on environmental uniqueness ^83^, i.e., if there are many positive sites having characteristics falling within given ranges of temperature, humidity, precipitation, etc., only one needs to be retained to represent this niche. We thinned using the function ′occfilt_env()′ from R package ′flexsdm′, v1.3.4 ^85^, based on nine purely environmental covariates (mean relative humidity, mean temperature, temperature range, temperature variation, total rainfall, max elevation, NDVI, distance to coast, distance to inland water), each stratified into six bins. Thinning was conducted independently for positive cells and pseudo-absence cells from each season’s training data from each dataset, resulting in between 188 and 1631 total cells (Table 3), excluding breeding season for period A which was not modelled due to lack of sufficient data.

**Table 3.**
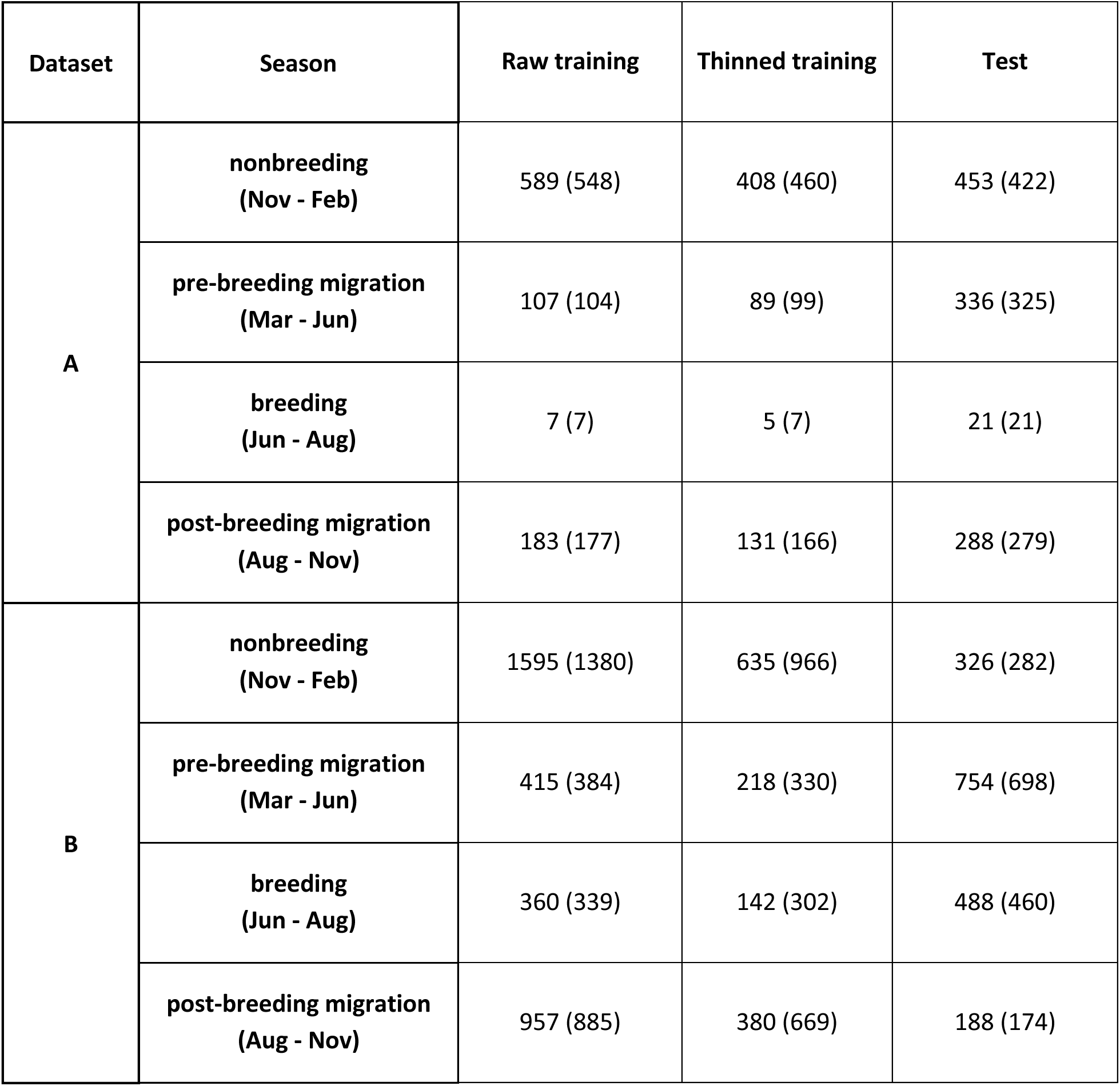
Data availability for model training and test sets in terms of number of grid cells. Value denotes number of positive grid cells, and bracketed value denotes number of grid cells sampled as pseudo-absences.

### Modelling approach

Using the above training data, we fitted BART, an ensemble-based nonparametric machine learning method based around constructing large numbers of decision trees ^86^. The decision tree structure of BART makes it ideal for modelling complex nonlinear relationships such as environment-host-disease interactions. An ensemble of candidate models is constructed via Markov Chain Monte Carlo (MCMC), with the predicted response value for a given covariate datapoint given by the mean predicted response across these candidate models. One particular advantage of BART is that credible intervals can be generated by taking quantiles of the ensemble of projected values. We carried out our model fitting using the R programming language package *embarcadero* ^87^, which offers an implementation of BART designed to interface with geospatial data and generate spatial projections; based on a fitted BART model *embarcadero* can generate a raster of predicted response variables from a multilayer raster of covariates.

We explore a basic model, plus a model including cross-seasonal covariates. In the basic model, we construct models using seasonally-variable environmental and ecological covariates for only the same behavioural season as the HPAI records being trained on, whereas with cross-seasonal covariates, we construct using seasonally-variable covariates from all four seasons. This is to allow for potential delay effects in the infectious disease dynamics; for instance, winter case numbers in a given location could be influenced by the local climate over summer, or large numbers of high-risk birds congregating in a location at one time of year could drive the circulation of infection lasting throughout the year. We also test a model formulation specifying country-level random intercepts (Supplemental Methods S1).

We performed five-fold cross validation to optimise the parameters of the MCMC, and performed stepwise variable set reduction to optimise our covariate set for predictive ability. We fit a separate instance of BART for each of our seasonal datasets, with the variable set reduction allowing models for different seasons to retain distinct covariates, reflecting annual variation in epidemiology. Uniform priors were specified for the probability of each covariate being selected at a given tree branching point, and on the exact split point of the covariate used in branching. A negative power law prior was specified for the probability of incrementing tree depth *d* as α(*1* + *d*)^−β^ where α was set to 2 and β was set to 0.95 following package defaults ^87^. We ran eight parallel MCMC chains of length 1000 plus a burn-in period of length 100, generating a total of 8000 posterior samples for each model instance.

### Comparison to domestic cases

We sense-checked our predictions by calculating the odds of observing a domestic bird HPAI detection in locations where domestic birds were present, stratified by model-projected risk. We extracted data on domestic bird detections of HPAI with a known H5 subtype from the same sources and over the same period as for wild bird cases. To ensure we only looked at cells with non-negligible domestic bird populations we added together chicken and domestic duck density raster layers to produce a raster capturing the combined density of both species. We first selected all 10km grid cells with > 1000 domestic ducks and chickens combined and labelled them as infected or uninfected based on the presence of at least one reported domestic case of HPAI. Each cell was also associated with a model-predicted probability of HPAI presence in wild birds. For each 10% band of predicted probability, we calculated the risk-stratified odds of infection as the number of cells with domestic bird H5 HPAI detection divided by the number of cells without (each restricted to those with > 1000 domestic ducks and chickens combined). For each period (A and B) and for each season this gave a set of 10 risk-stratified odds which were visualised to examine potential linkage between HPAI outbreaks in wild and domestic birds.

The code used to generate our training data and perform our analysis is available at https://github.com/sarahhayes/avian_flu_sdm.

## Results

Based on performance metrics in predicting held-out test data (Table 4), our BART models trained on environmental and ecological predictors showed good generalisability in understanding the distribution of H5 HPAI clade 2.3.4.4b for each modelled season in both data periods (Table 4). Specificity and sensitivity were generally well-balanced, though for the ongoing (since 2021) H5N1 outbreak performance in warmer periods (pre-breeding migration and breeding seasons) was slightly weaker in sensitivity and overall Area Under Receiver-Operating Characteristic curve (AUROC), likely due to much smaller training data availability compared to test set for these seasons (Table 3; Supplemental Figure S1B). Including a random intercept term stratified by country did not noticeably improve overall AUROC for either dataset and reduced sensitivity in some cases (Supplemental Methods S1; Supplemental Table S1). Allowing models to access cross-seasonal covariates had surprisingly minimal impact upon model performance (Table 4), though we retain these as our preferred model suite to allow us to investigate ecological impacts of annual migration cycles on year-round avian influenza risk.

**Table 4.** Posterior means of performance metrics for each model option on each dataset’s test set. AUROC denotes Area Under the Receiver-Operating Characteristic curve, CV denotes covariates.

|  |  |  | Model |  |
| --- | --- | --- | --- | --- |
|  |  |  | base model | + cross-seasonal CV's |
| A | nonbreeding (Nov - Feb) | Sens.<br>Spec.<br>AUROC | 0.77 (0.76; 0.72-0.81)<br>0.83 (0.82; 0.77-0.87)<br>0.90 (0.88; 0.85-0.90) | 0.80 (0.80; 0.76-0.84)<br>0.81 (0.80; 0.75-0.85)<br>0.90 (0.88; 0.86-0.90) |
|  | pre-breeding migration (Mar - Jun) | Sens.<br>Spec.<br>AUROC | 0.77 (0.77; 0.69-0.83)<br>0.89 (0.83; 0.74-0.90)<br>0.91 (0.88; 0.84-0.90) | 0.86 (0.83; 0.76-0.88)<br>0.83 (0.77; 0.65-0.86)<br>0.92 (0.89; 0.85-0.91) |
|  | breeding (Jun - Aug) | Sens.<br>Spec.<br>AUROC | - | - |
|  | post-breeding migration (Aug - Nov) | Sens.<br>Spec.<br>AUROC | 0.88 (0.86; 0.77-0.91)<br>0.79 (0.76; 0.64-0.83)<br>0.86 (0.85; 0.80-0.90) | 0.86 (0.86; 0.80-0.91)<br>0.75 (0.74; 0.68-0.78)<br>0.86 (0.85; 0.81-0.89) |
| B | nonbreeding (Nov - Feb) | Sens.<br>Spec.<br>AUROC | 0.83 (0.81; 0.77-0.84)<br>0.84 (0.84; 0.80-0.90)<br>0.93 (0.91; 0.89-0.93) | 0.80 (0.80; 0.77-0.83)<br>0.83 (0.83; 0.80-0.87)<br>0.91 (0.90; 0.88-0.92) |
|  | pre-breeding migration (Mar - Jun) | Sens.<br>Spec.<br>AUROC | 0.80 (0.78; 0.72-0.84)<br>0.61 (0.61; 0.53-0.69)<br>0.79 (0.75; 0.71-0.79) | 0.77 (0.75; 0.70-0.80)<br>0.69 (0.64; 0.56-0.71)<br>0.79 (0.75; 0.71-0.79) |
|  | breeding (Jun - Aug) | Sens.<br>Spec.<br>AUROC | 0.84 (0.81; 0.76-0.85)<br>0.65 (0.62; 0.56-0.68)<br>0.79 (0.77; 0.74-0.80) | 0.83 (0.81; 0.77-0.84)<br>0.64 (0.62; 0.57-0.67)<br>0.76 (0.76; 0.72-0.79) |
|  | post-breeding migration (Aug - Nov) | Sens.<br>Spec.<br>AUROC | 0.83 (0.80; 0.75-0.85)<br>0.84 (0.78; 0.67-0.84)<br>0.88 (0.86; 0.81-0.88) | 0.80 (0.77; 0.72-0.83)<br>0.85 (0.83; 0.77-0.87)<br>0.89 (0.87; 0.85-0.89) |

We generated 10km resolution risk projections for HPAI clade 2.3.4.4b across Europe with 95% credible intervals (Figure 2). Considering discrete outbreaks of H5N6 and H5N8 clade 2.3.4.4b (period A), risk hotspots for HPAI in wild birds were present across central Europe during the aggregated non-breeding season, dissipating northward during pre-breeding migration (Figure 2A). While the breeding period lacked sufficient data to model, we note a more limited risk distribution during post-breeding migration (Figure 2A). Restricting to highest-confidence hotspots of HPAI presence (high risk at 2.5th percentiles) showed consistent risk concentrated around central and eastern European shorelines and waterways, while restricting to highest confidence hotspots of HPAI absence (low risk at 97.5th percentiles) suggested low suitability for HPAI in high-elevation, mountainous regions as well as the Iberian peninsula.

**Figure 2.**
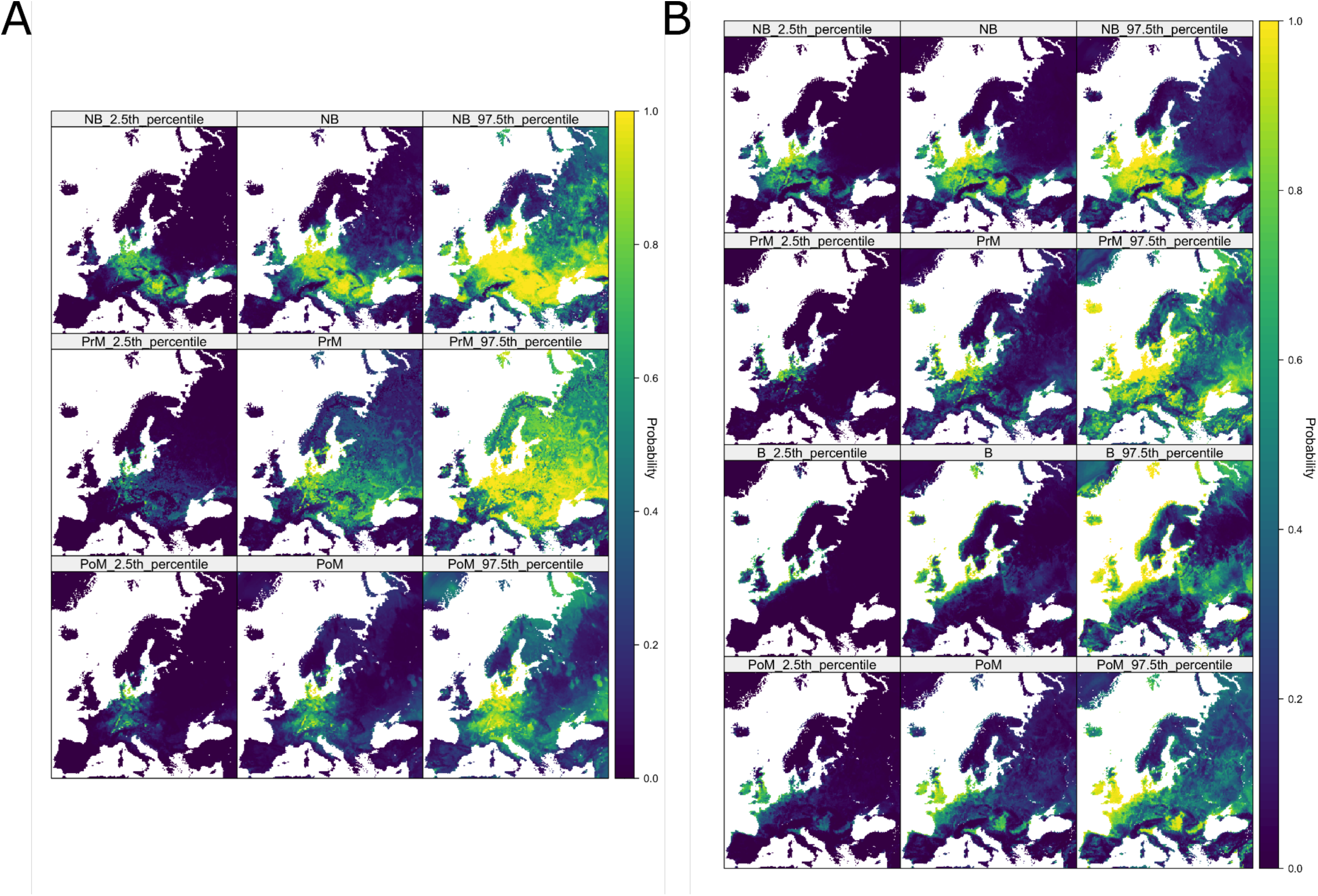
Projected risk maps for H5 HPAI clade 2.3.4.4b and associated uncertainty intervals. Projected probability of H5 HPAI clade 2.3.4.4b presence by aggregated bird behavioural season (NB = Nonbreeding season, PrM = Pre-breeding migration, B = Breeding season, PoM = Post-breeding migration) during period A (10/8/2016 to 9/8/2021, spanning H5N8 and H5N6 events) and period B (10/8/2021 to 29/2/2024, ongoing H5N1 outbreak) from selected final models with cross-seasonal covariates. Central columns are posterior means from BART ensemble, left and right columns represent 95% credible intervals.

Contrastingly, we found the ongoing H5N1 epizootic in wild birds (period B) to have much more spatially concentrated risk (Figure 2B). Predicted H5N1 HPAI hotspots appeared further westward and centred around the British Isles and North Sea coast with comparatively lower risk throughout eastern Europe (Figure 2B). Presence of H5N1 also appeared less seasonally-variable than for H5N6 and H5N8, with only minimal northward shift during pre-breeding migration (Figure 2B), though risk during the breeding season was highly localised to north-western European coastlines. Although different environments were observed to be at higher risk compared to previous outbreaks, the lowest risk was again consistently observed for more mountainous and arid regions.

Presence data in test set A is from the 2020-2021 H5N8 HPAI clade 2.3.4.4b outbreak. Presence data in test set B is from 1/3/2023 to 29/2/2024, during the ongoing (since 2021) H5N1 HPAI clade 2.3.4.4b outbreak. Values in brackets denote posterior median and 95% credible interval of metrics within individual trees. As posterior mean values are based on consensus predictions averaging over all sampled trees, this can exceed posterior medians and 95% credible intervals.

These continental-scale differences can be explained by differing importance of bioclimatic, topographic, livestock demographic, and host ecological factors in dynamics of respective HPAI outbreaks (Figure 3, Figure 4). During H5N8 and H5N6 events of HPAI clade 2.3.4.4b, bioclimatic, topographic, and host ecological variables were each well-represented after variable selection in models of each season (Figure 3A). The presence of lagged variables among the bioclimatic variables retained for period A models pointed to an association between colder months (within nonbreeding season; November to February) and risk. Partial dependence curves showed that more variable temperatures over the respective season led to greater predicted risk of influenza (Figure 4A.i), and that lower zero-degree isotherms (i.e., freezing temperatures nearer ground level) were associated with risk in nonbreeding and post-breeding migratory seasons (Supplemental Figures S3, S5). No overall topographic patterns dominated, with different variables being retained for different seasons, e.g., savanna (defined as land with 10-30% cover of tree canopies >2m) was associated with decreased risk for pre-breeding migration (Figure 4A.ii) while low-lying, flat land was associated with increased risk for the nonbreeding season (Supplemental Figure S4).

**Figure 3.**
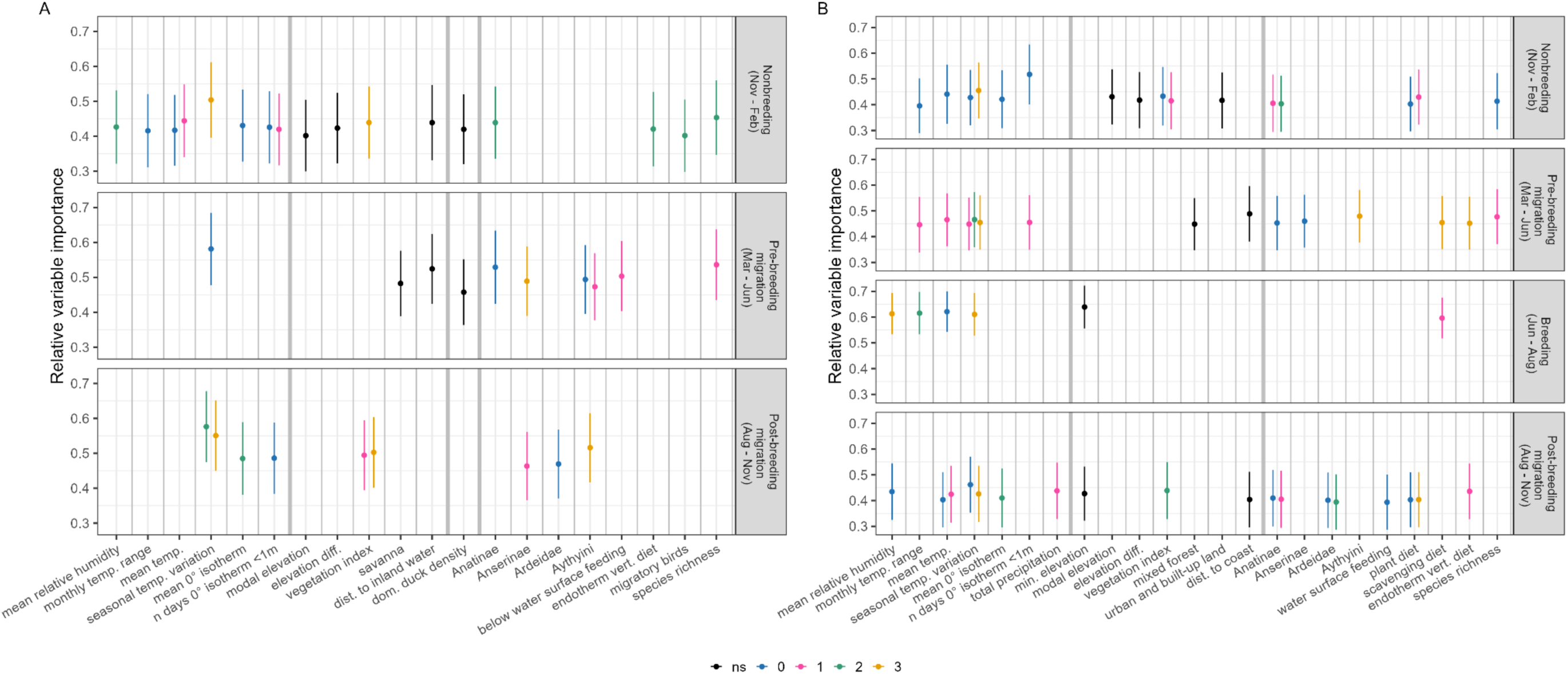
Variable importance of BART models. Relative variable importance of each covariate kept in optimal fitted BART models for period A (10/8/2016 to 9/8/2021, spanning H5N8 and H5N6 events) and period B (10/8/2021 to 29/2/2024, ongoing H5N1 outbreak). Points denote normalised proportion of times variable was used in a tree decision split out of all splits, averaged over 8000 draws from the posterior tree space. Error bars denote +/- 1 SD. Colour key denotes precedence of bird behavioural season of covariate to modelled season, either, 0 (same season, e.g., breeding season model using breeding season covariate) or 1 (one season prior, e.g., breeding season model using pre-breeding migration season covariate), 2, or 3 season prior; black denotes non-seasonally-variable covariates (“ns”). Thicker grey lines separate bioclimatic, topographic, livestock demography, and host ecological covariates.

**Figure 4.**
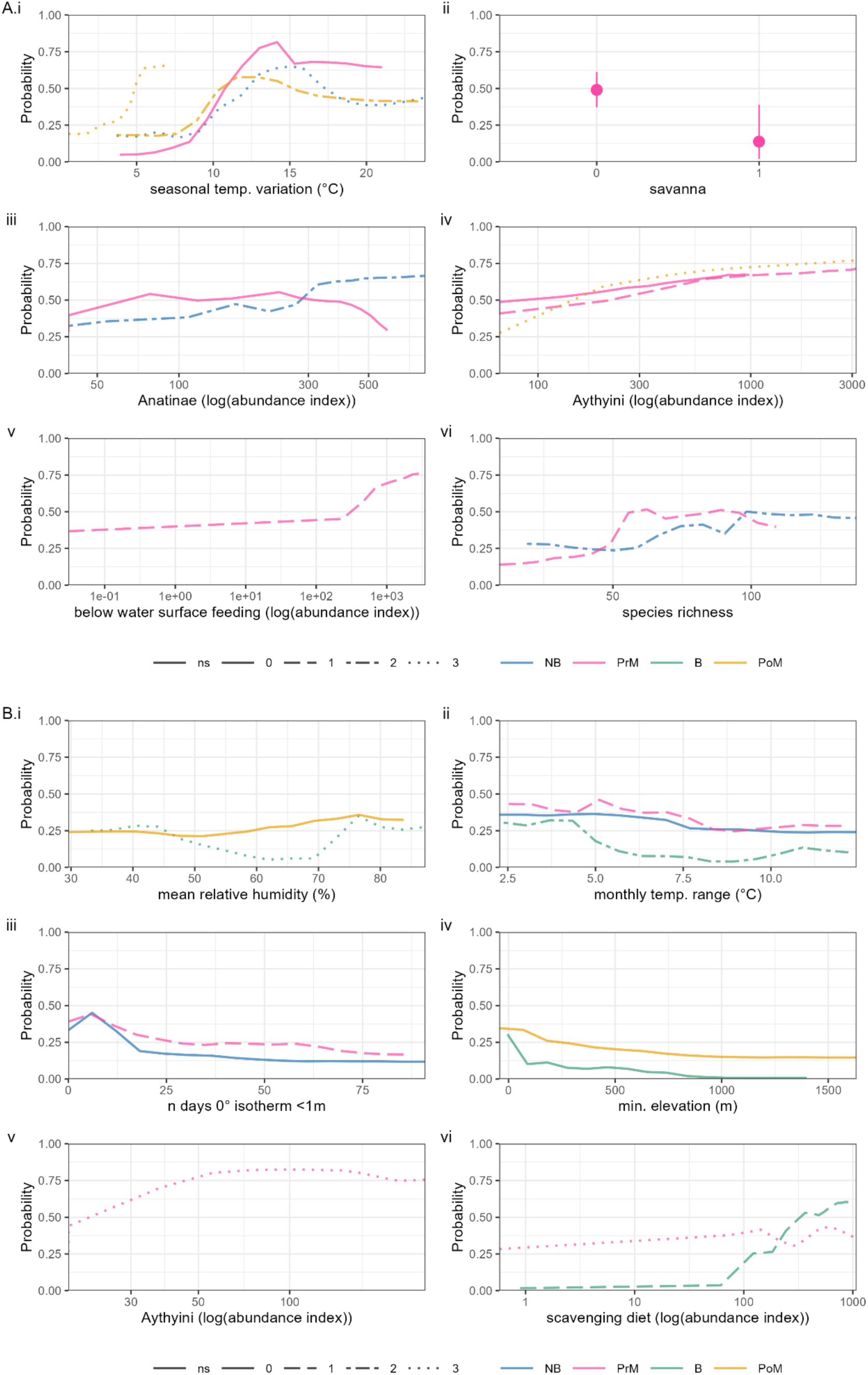
Partial dependence of BART models. Partial dependence associated with top six covariates by average variable importance in final fitted BART models for period A (10/8/2016 to 9/8/2021, spanning H5N8 and H5N6 events) and period B (10/8/2021 to 29/2/2024, ongoing H5N1 outbreak). Y axis denotes marginal probability of HPAI presence, i.e., averaging out effects of all other covariates. Solid lines denote median values over 8000 draws from the posterior tree space. Colour denotes separate modelled bird behavioural seasons (NB = Nonbreeding season, PrM = Pre-breeding migration, B = Breeding season, PoM = Post-breeding migration) of HPAI and line type denotes seasonal delay increasing from 0 (solid) to 3 (dotted) seasons prior.

Each season’s model in period A also emphasised the importance of host ecology, particularly the pre-breeding migration model (Figure 3A). Many ecological variables retained related to population sizes during the breeding season via lagged effects, suggesting predictable annual cycling in influenza risk. Specifically, abundance indices of Anatinae, Aythyini, and to a lesser extent, Anserinae and Ardeidae showed generally consistent increases in risk with increasing host population (Figure 4A.iii, iv; Supplemental Figures S3-S5). Behavioural and dietary traits showed minor, season-dependent effects, e.g., risk association with increasing population of underwater feeders (Figure 4A.v) or predatory birds (Supplemental Figure S3). Notably, the collective population of migratory species during the breeding season was predictive of risk during the subsequent non-breeding season (Supplemental Figure S3). Risk of HPAI also increased with a more diverse host community in species richness, up to a plateau at ∼50-100 species (Figure 4A.vi).

Models trained on geolocated H5N1 HPAI clade 2.3.4.4b data in period B also featured a combination of bioclimatic, topographic, and ecological variables with some key differences. As in period A, winter climate, including humidity, temperature, temperature variation, and zero-degree isotherms of post-breeding and nonbreeding seasons were consistently associated with HPAI presence throughout the year (Figure 3B, Figure 4B.i - iii). For topography, low elevation was again associated with greater risk (Figure 4B.iv). Compared to period A, however, land cover had additional consistent associations with influenza risk, including greater risk in denser vegetated areas, urban and built-up areas, and coastal areas (Supplemental Figures S6, S7, S9), consistent with topography of the projected risk maps.

Among potential high-risk host taxa, abundance index of *Aythyini* had the highest average importance in period B models (Figure 3B), associated with a relatively rapid increase and plateau in predicted influenza risk during the pre-breeding migratory season (Figure 4B.v). Other taxa had predictive effects with different annual lags e.g., breeding populations of *Anatidae* and migratory populations of *Ardeidae* and *Anserinae* affecting later seasons (Supplemental Figures S6, S7, S9). Dietary and behavioural species-trait abundance indices were retained across all seasons, with partial dependence curves tending to capture sharp risk increases beyond threshold populations, e.g., of scavenging birds (Figure 3B.vi, Supplemental Figures S6 - S9). A positive risk relationship was again retained with species richness, though only richness during the nonbreeding season was influential (Supplemental Figures S6, S7).

Overall, the season-specific effects of these traits suggests that localised ecology (i.e., small-scale interactions between different covariates) is important to understanding the distribution of H5 clade 2.3.4.4b influenza.

Our comparison of model predictions with domestic H5 HPAI detections indicated a trend towards increased odds of infection in domestic birds with model-projected risk of HPAI presence in wild birds for both model periods (Figure 5). Spatial concordance between model-generated predicted probabilities and reports of HPAI in domestic birds is visualised in Supplemental Figure S10.

**Figure 5:**
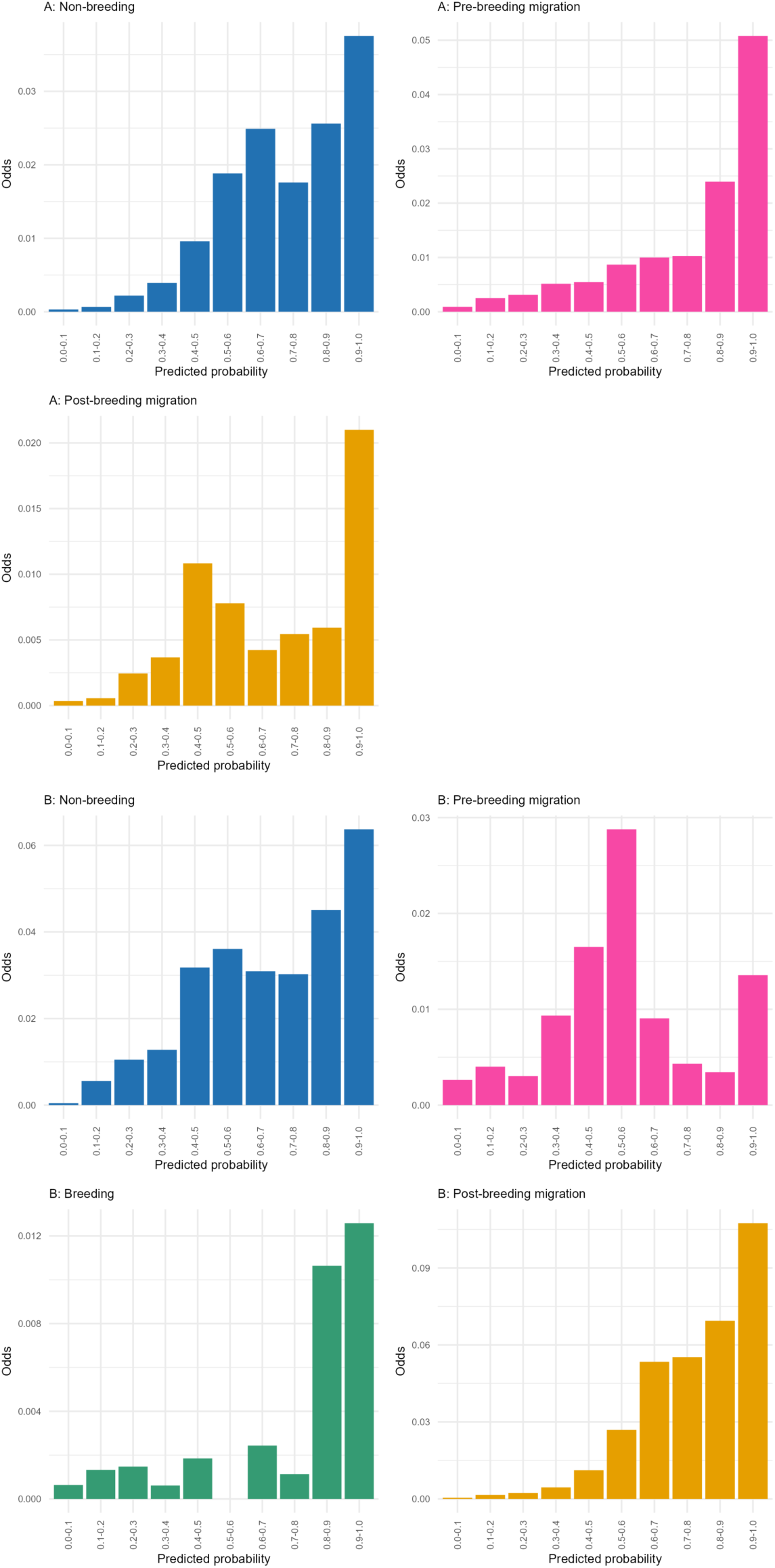
Comparison of odds of reporting H5 clade 2.3.3.4b HPAI in domestic birds stratified by model-predicted risk in wild birds. Odds of H5 infection in domestic birds for each period and season by model predicted probabilities of H5 clade 2.3.4.4b HPAI risk for wild birds.

## Discussion

In this study we have demonstrated how inclusion of both environmental and ecological covariates can lead to generalisable models of avian influenza distribution in wild birds, particularly in colder months of the year. Our analysis reveals subtle differences in the epidemiology of clade 2.3.4.4b of HPAI once H5N1 became the dominant subtype within Europe. This is reflected both in the projected geospatial risk distributions, and in which covariates best explain these distributions between period A and period B models. Our projections suggest that the ongoing outbreak of the H5N1 genotype is associated with an increased risk of avian influenza presence around northwest Europe, in agreement with the observed large numbers of HPAI detections in this region. Underlying risk patterns were more ambiguous in eastern Europe due to less intensive sampling, though our projections suggest that while H5N8 and H5N6 HPAI widely dispersed across these areas during non-breeding and migratory seasons, the incursion of H5N1 was limited to waterways and shores.

In line with these observed patterns of detection, we expect risk to be associated with proximity to coastlines, low elevations, low temperatures, and presence of high-risk waterbirds. We found that environmental factors reflecting physical geography and climate tended to be more consistent than wild bird ecology in predicting HPAI risk, possibly reflecting inherent constraints on wild bird ecology imposed by background environmental factors. Our results suggest that while risk is indeed highest in the heavily sampled regions of northwest Europe, there is also a strong potential for HPAI presence in regions where case detections are less frequent. From a theoretical perspective, the incorporation of ecological risk factors into HPAI presence/absence models appears to offer a small but distinct improvement in predictive ability.

The minimal differences in performance when adding country-level random intercepts suggest that any spatial heterogeneity in our detection data was effectively captured by weighting for background citizen surveillance of birds in our pseudo-absence generation process. However, the very low numbers of detections in specific eastern European countries (e.g., Albania, Belarus, Moldova, Ukraine) suggest that there are indeed national-level differences in detection and reporting rates which are difficult to estimate due to the absence of information from these countries in many of our training datasets. The selection of many cross-seasonal covariates during model fitting suggests that future mechanistic models may need to incorporate delay effects to fully capture the complex dynamics of avian influenza outbreaks in wild birds.

Our risk projections point to an increased risk of HPAI presence in northwest Europe, particularly around the North Sea coast, associated with the emergence of the H5N1 genotype of clade 2.3.4.4b. The confidence limits for our projections suggest that the Black Sea coast may also be a zone of high H5N1 risk, as it was for the previous H5N8 subtype of clade 2.3.4.4b. This shift in geographical burden of risk appears to be driven by differing importance of climate, geography, and abundance of specific bird taxa (*Anatinae*, *Anserinae*, *Ardeidae*, and *Aythyini*) at different calendar times and with different seasonal lags.

SDMs trained on clades prior to 2.3.4.4b have established a consistent link between temperature and avian influenza in wild birds ^48,49,51,52^, with further study highlighting the association between both temperature-driven waterbird movements and congregation of waterbirds at the zero-degree isotherm and the distribution and spread of HPAI ^88^. We report this association with temperature and zero-degree isotherm variables also holds for H5 clade 2.3.4.4b in our models. At least one temperature variable was selected for all models and temperature variation had high variable importance across both periods A and B, whilst zero-degree isotherm variables were selected for all bird behavioural seasons except pre-breeding migration in period A and the breeding season in period B. The suggested influence of temperature on current HPAI risk in wild birds is likely to be multifactorial and may reflect features such as habitat suitability for high-risk hosts, migratory bird movements, congregation of birds near unfrozen water sources during winter months and duration of survival of virus in the environment ^30,31,88^. The importance of temperature in our models also suggests that anthropogenic climate change is likely to drive changes in the spatiotemporal distribution of avian influenza in Europe. Relatedly, we find minor, but detectable effects of land cover type on HPAI risk, with lower average risk in savanna-type or mixed-forest biomes and greater average risk in urban or built-up lands. This may hint at a role of anthropogenic habitat disruption and fragmentation upon influenza epidemiology, as empirically shown for many other zoonotic disease systems ^89^.

By conducting variable selection we demonstrate that incorporating ecology provides a unique improvement in models of avian influenza presence; all models featured ecological covariates, with most combining a range of different bird population indices. This identifies wild bird ecology as a distinct source of information not present in background environmental data. In contrast to the more consistent set of climate covariates, ecological covariates varied seasonally with exact indicator taxa and/or trait abundance indices preferentially being retained in models of different bird behaviours and with different temporal lags. This suggests that avian influenza incidence is primarily driven by patterns of host species-level climatic and environmental suitability but is also subject to highly localised ecological factors for H5 clade 2.3.4.4b.

We observed an association with bird species richness for nonbreeding and pre-breeding migratory seasons. Whether host diversity generally increases or decreases disease risk is somewhat contested ^90^ and depends on scale, pathogen traits, and host species composition. We report a positive relationship between species richness and HPAI presence, potentially capturing greater opportunities for contact and transmission between different populations ^28^ and/or importance of additional key host taxa following Huang *et al.* reporting a similar positive effect of richness of “higher-risk hosts” upon HPAI in Europe from 2005-2008 ^50^.

Among host taxa we explicitly modelled, relative abundance of *Anatinae* (dabbling ducks) was the most consistent predictor of HPAI presence, followed by *Aythini* (pochards), *Anserinae* (swans and geese), and *Ardeidae* (herons). Except for *Ardeidae*, these taxa are generally considered maintenance hosts of avian influenza as part of the Anseriformes. Aside from topographic variables like elevation and distance to wetland, the most informative predictors of avian influenza in SDMs for Japan and South Korea were population measures or correlates of *Anatidae* ^53,91^. Mallards within the *Anatidae* appear to be effective carriers as they may experience only mild disease, even when infected with H5N1 clade 2.3.4.4b ^25^. In phylogenetic discrete trait analyses of the North American H5N1 clade 2.3.4.4b outbreak, longer persistence of infection and high rates of transmission to other taxa were observed for Anseriformes, as well as a Charadriiformes ^22^. Despite this, we did not detect any association between abundance of Charadriiformes hosts (*Laridae* (gulls), *Arenaria* (turnstones), or *Calidris* (sandpipers)) and HPAI presence. This suggests that while gulls and shorebirds have experienced unprecedented infection and die-offs, they may not have driven the precise spatial dynamics of HPAI clade 2.3.4.4b in Europe.

Although the total abundance index of migratory birds was only retained in a single model of period A, the effects of migration may be reflected in the changing predictive power of these taxa throughout the calendar year.

Alongside the abundance estimates for high-risk taxa, our construction of species-trait abundance indices represents a novel attempt to quantify the effect of ecological factors on HPAI risk at the continental scale. Although they improved models, overall abundance indices of birds with specific diets and foraging behaviours showed varying and highly localised trends in risk (Supplemental Figures S3 - S9). Foraging behaviour has previously been suggested as a major driver of avian influenza ^28^, as faecal-oral transmission primarily occurs through viral shedding into water. However, studies looking at viral shedding of HPAI including clade 2.3.4.4b viruses have reported high levels of additional shedding via the respiratory system ^32–35^. We find a somewhat unexpected importance of scavenging and predation behaviour in both modelled periods. Combined with the high reported rates of cross-species transmission to predatory birds such as owls and raptors ^22^, this indicates the role of predation in spatial transmission should not be ignored.

Equivalent functional ecology and life history measures have been found to act as predictors of individual-level infection with previous clades ^50,92^, and emerging studies are highlighting similar trends for community-level infection, where plant consumption was predictive of HPAI clade 2.3.4.4b infection ^27^, consistent with our findings for H5N1 in period B. The relatively low importance assigned to functional ecological factors in our models is potentially due to a degree of redundancy in the models with both taxon abundance layers and species-trait abundance layers; the derived nature of the species-trait abundance layers means that they carry an inherently higher level of error than the taxon abundance layers, while contributing a limited amount of independent information compared to the taxon abundance layers. For instance, knowing the spatial abundance of specific high-risk waterfowl taxa may contribute the same information as knowing the combined species-trait abundance layers for plant-feeding, feeding around the water surface, and feeding below the water surface.

Continental-scale modelling efforts to understand avian infectious diseases are critically necessary, given their ability to disseminate long distances through bird migration ^26^; recent phylodynamic models estimate it took only five months for a single introduction of 2.3.4.4b HPAI to North America to spread from east coast to west coast via migratory flyways ^22^. However, if we are to better capture host dynamics at these scales and improve spatial projections of avian influenza, we need higher-quality underlying ecological data. The spatial ecology layers we use come with several important caveats. In the absence of comprehensive survey data, we use projected abundance layers from eBird Status and Trends products and global species population size estimates. Our models are therefore effectively downstream from another set of models and thus need to be considered in light of any uncertainties and limitations associated with those models ^93^. The projections from eBird Status and Trends also only covered a subset of all bird species known to be present in Europe, meaning that our ecological covariates could potentially underestimate abundances in spatially heterogeneous ways. The role of ecological covariates in HPAI risk prediction points to high-confidence models of wild bird abundance for a more comprehensive selection of species and covering a wider geographical area as a key target for future research. Automated “computer vision” systems to recognise presence, abundance, and location of species from digital images and their metadata could prove useful in this context ^94^.

There are also substantial challenges associated with working with current standards of HPAI case reports. Although HPAI is classified as a listed disease by the WOAH meaning that national authorities are required to report HPAI viruses detected in both poultry and non-poultry species, including wild birds ^95^, surveillance strategies across countries are not standardised and may differ substantially between countries in the same region ^96,97^. This results in significant underreporting of wild bird HPAI, with gaps increasingly evident in the wake of the expansive wild bird die-off associated with clade 2.3.4.4b ^98^. Passive surveillance is frequently used and relies on submission of sick or dead wild birds. Rates of data submission are likely to show spatial and temporal variation based on factors such as human population density and public awareness of disease, whilst testing capacity may also vary over time depending on factors such as disease priority and demand. Similar variation also exists across active surveillance strategies. In this study we responded to these data quality issues by considering presence/pseudo-absence at the geographic level rather than finer-grained reported incidence data, which are associated with inherently higher levels of error due to the more complex nature of the signal being measured. Consistency in surveillance both within and between countries, reporting of negative tests and/or submission rates from passive surveillance, and accurate and consistent labelling of affected species would significantly improve data interpretability for end users. Higher quality of reporting data should improve the predictive ability of complex mechanistic models, as well as broader presence/absence approaches as in our work.

In this study we have identified a shift in wild bird HPAI risk towards northwest Europe following the dominance of the H5N1 genotype of clade 2.3.4.4b, which is associated with changes to the underlying predictive factors driving HPAI epidemiology. Our work also serves as an assessment of the role of ecological factors in predictive models of HPAI, with our results suggesting that these factors can improve model performance, particularly if more accurate quantitative measures of wild bird ecology become available in the future.

## Acknowledgements

We thank Sarah Hill, Stephen Vickers, Kieran Sharkey, Stephen Brough, Michelle Wille, and John Drake for their helpful comments on this work. This work was undertaken on Barkla, part of the High Performance Computing facilities at the University of Liverpool, UK. JH, MB, JM, and LB would like to acknowledge funding support by CSL Seqirus UK Ltd to the University of Liverpool through the 2022 – 2027 Framework Research Collaboration Agreement Influenza Research Partnership, collaboration plan 002. SH and CAD would like to acknowledge funding support from the National Institute for Health and Care Research Unit Emerging and Zoonotic Infections (Grant HPRU200907). CAD would also like to acknowledge funding support from the Medical Research Council (MRC) Centre for Global Infectious Disease Analysis (grant number MR/R015600/1), which is jointly funded by the UK MRC and the UK Foreign, Commonwealth and Development Office (FCDO), under the MRC/FCDO Concordat agreement and is also part of the EDCTP2 programme supported by the European Union (EU).

## Supplementary information

**Supplemental Figure S1.**
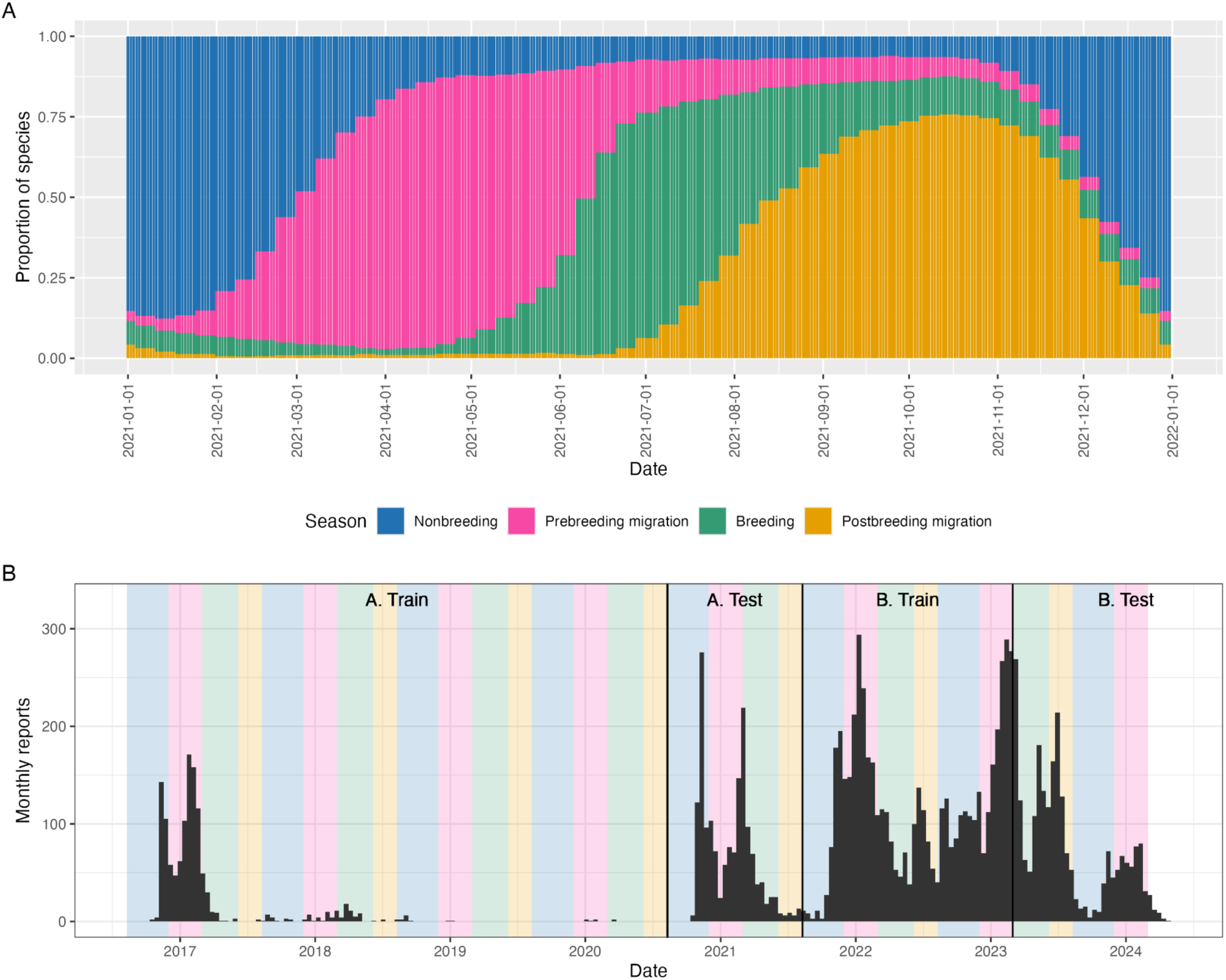
Aggregated Europe-wide bird behavioural seasons. A) Distribution of behavioural season of seasonal European bird species over the course of the calendar year 2021, based on eBird Status and Trends records. B) Total independent incident reports of HPAI clade 2.3.4.4b in wild birds during model training/test periods, with shaded time strips denoting annual seasonal periods as chosen considering panel A, demonstrating the well-established persistence during warmer breeding and post-breeding migration seasons associated with H5N1 HPAI during period B.

**Supplemental Figure S2.**
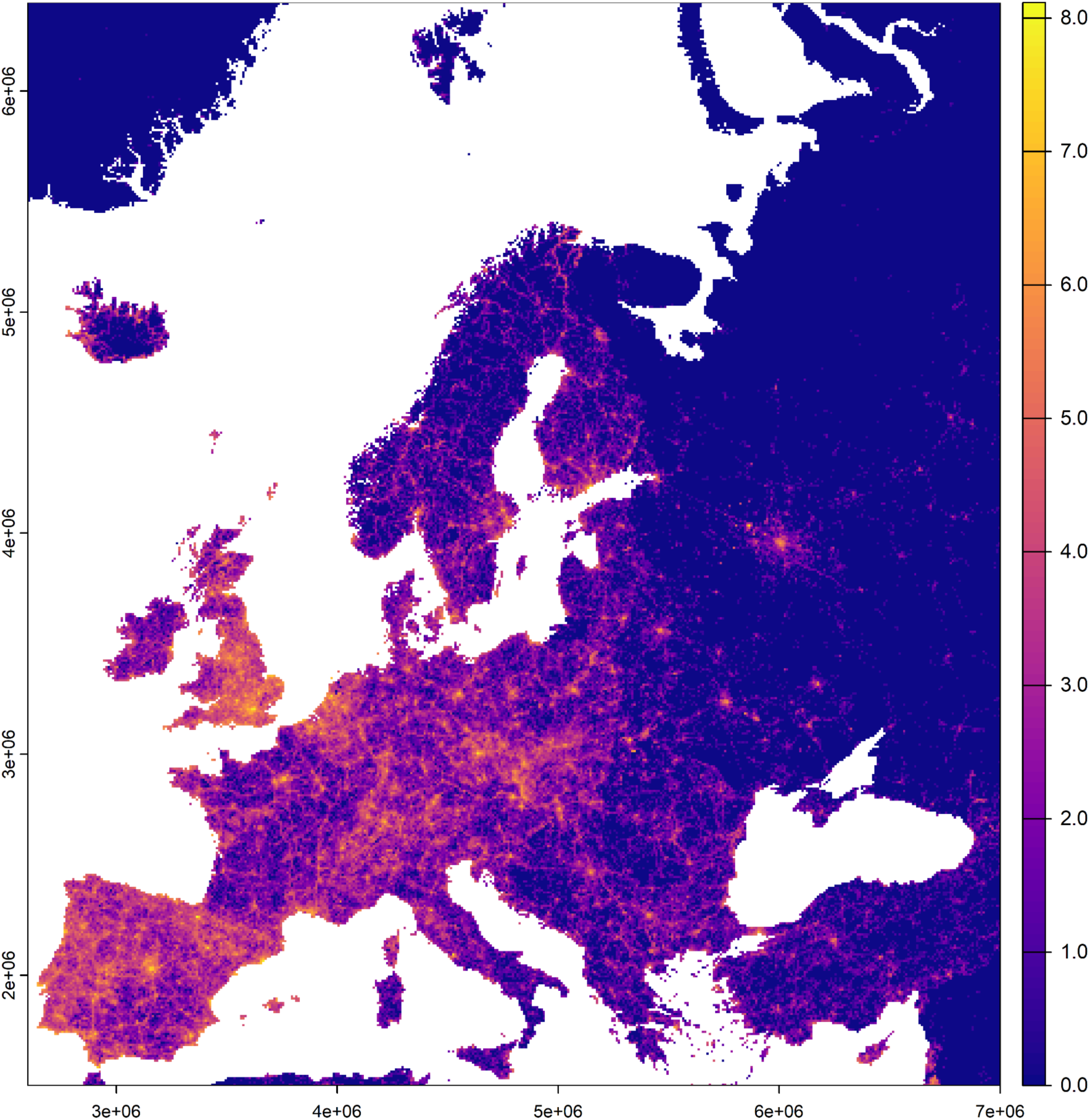
Mapped density of eBird sightings. Mapped density of bird sighting records in Europe from the eBird citizen surveillance platform. Log-transformed total sightings were counted as instances of unique data-user-geolocations over duration of study period (10/8/2016 - 29/2/2024).

**Supplemental Figure S3.**
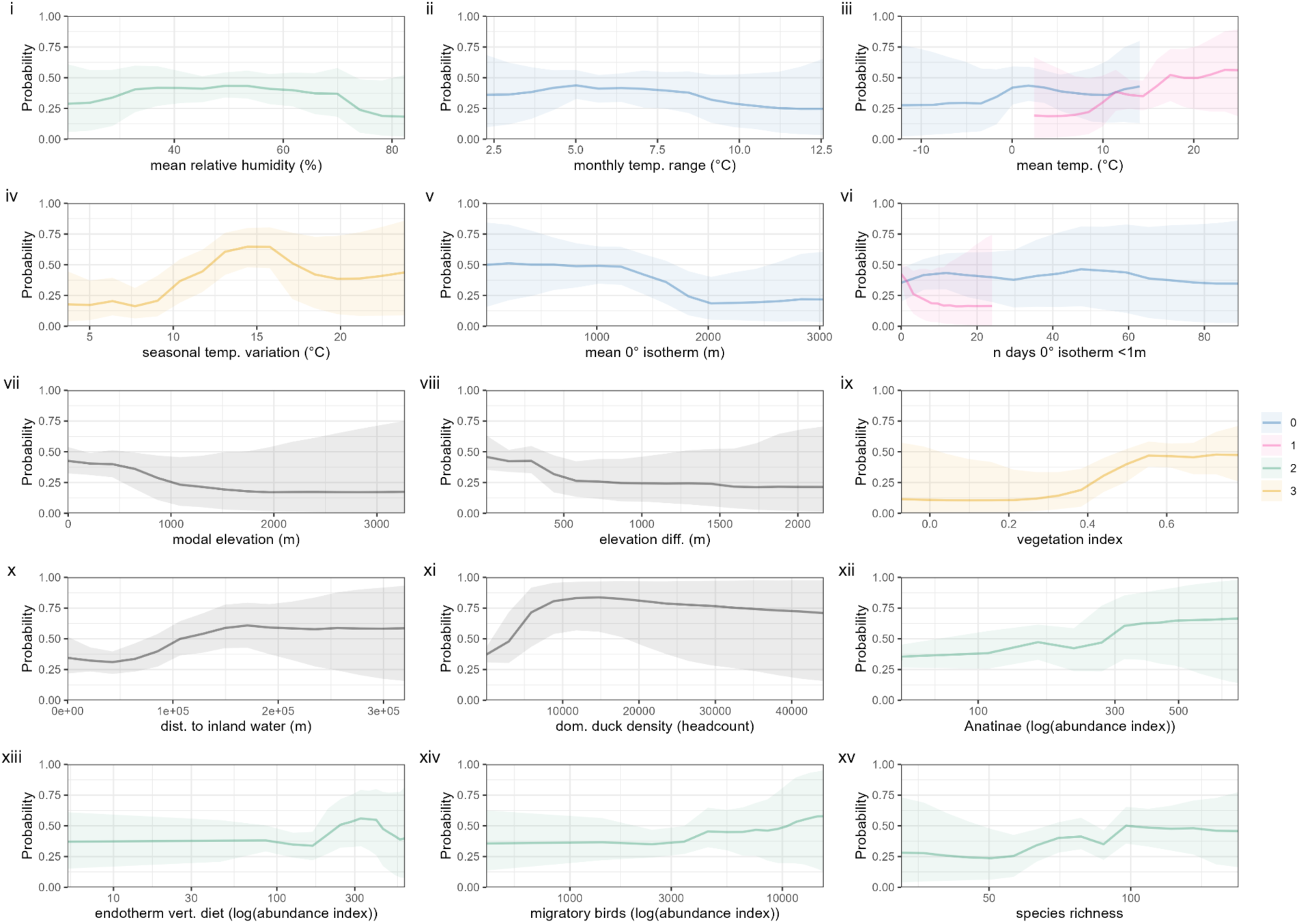
Full partial dependence of BART model trained on dataset A, non-breeding season. Partial dependence associated with all covariates in final BART model of aggregated non-breeding season (30th November - 28th February) fitted to geospatial HPAI data for period A (10/8/2016 to 9/8/2021, spanning H5N8 and H5N6 events). Y axis denotes marginal probability of HPAI presence, i.e., averaging out effects of all other covariates. For continuous covariates, solid lines denote median values while shaded areas denote uncertainty via 2.5th percentile and 97.5th percentile values over 8000 draws from the posterior tree space. For categorical covariates, points denote median values while error bars denote via 2.5th percentile and 97.5th percentile values. Colours denote seasonal delay increasing from 0 (predicted season) to 3 (three seasons prior to predicted season); black denotes non-seasonally-variable covariates.

**Supplemental Figure S4.**
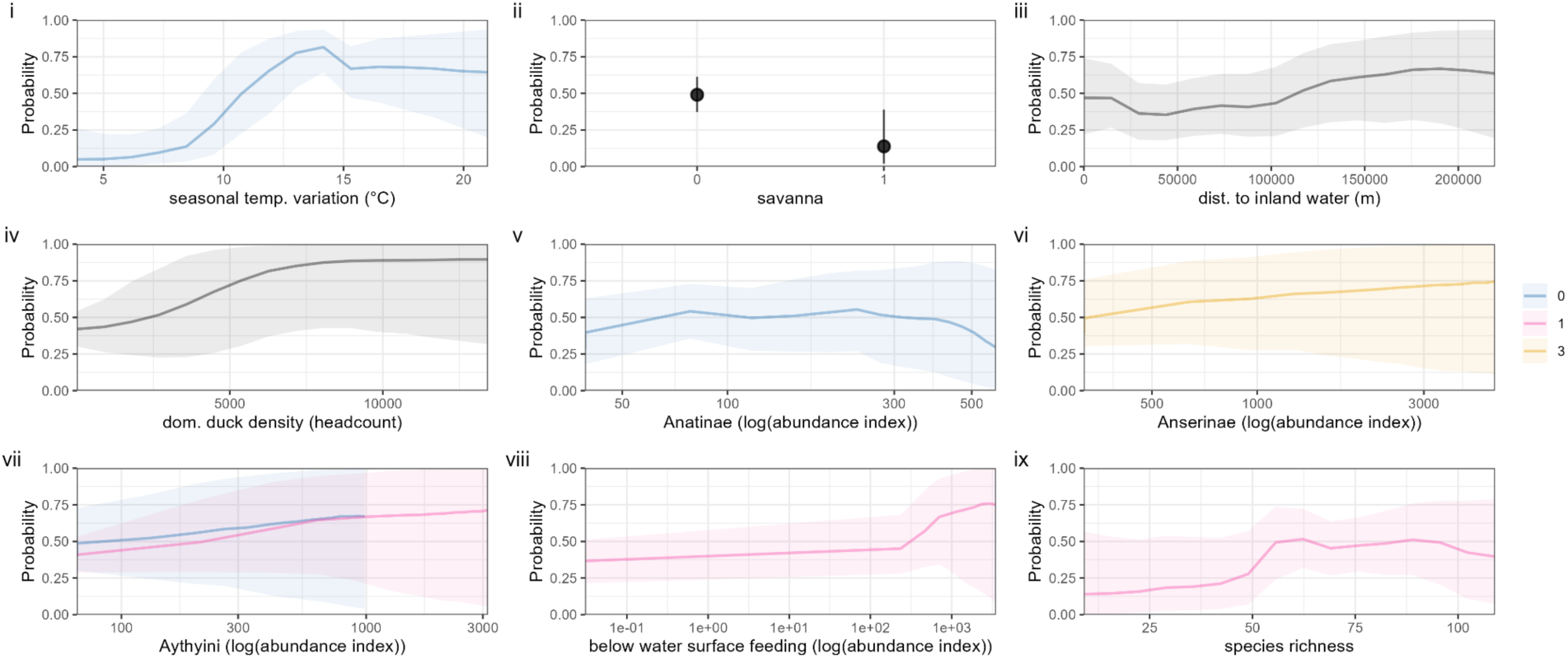
Full partial dependence of BART model trained on dataset A, pre-breeding migration. Partial dependence associated with all covariates in final BART model of aggregated pre-breeding migration season (1st March - 6th June) fitted to geospatial HPAI data for period A (10/8/2016 to 9/8/2021, spanning H5N8 and H5N6 events). Y axis denotes marginal probability of HPAI presence, i.e., averaging out effects of all other covariates. For continuous covariates, solid lines denote median values while shaded areas denote uncertainty via 2.5th percentile and 97.5th percentile values over 8000 draws from the posterior tree space. For categorical covariates, points denote median values while error bars denote via 2.5th percentile and 97.5th percentile values. Colours denote seasonal delay increasing from 0 (predicted season) to 3 (three seasons prior to predicted season); black denotes non-seasonally-variable covariates.

**Supplemental Figure S5.**
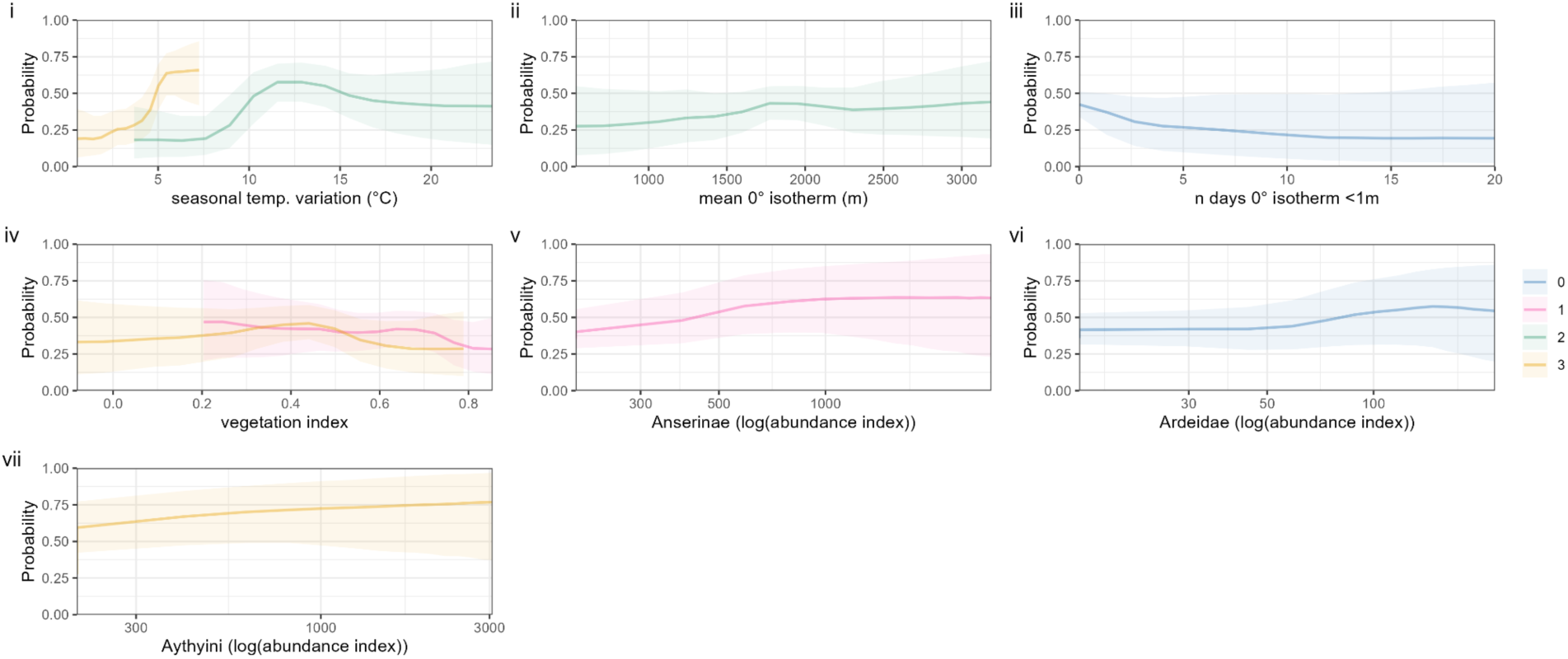
Full partial dependence of BART model trained on dataset A, post-breeding migration. Partial dependence associated with all covariates in final BART model of aggregated post-breeding migration season (10th August - 29th November) fitted to geospatial HPAI data for period A (10/8/2016 to 9/8/2021, spanning H5N8 and H5N6 events). Y axis denotes marginal probability of HPAI presence, i.e., averaging out effects of all other covariates. For continuous covariates, solid lines denote median values while shaded areas denote uncertainty via 2.5th percentile and 97.5th percentile values over 8000 draws from the posterior tree space. For categorical covariates, points denote median values while error bars denote via 2.5th percentile and 97.5th percentile values. Colours denote seasonal delay increasing from 0 (predicted season) to 3 (three seasons prior to predicted season); black denotes non-seasonally-variable covariates.

**Supplemental Figure S6.**
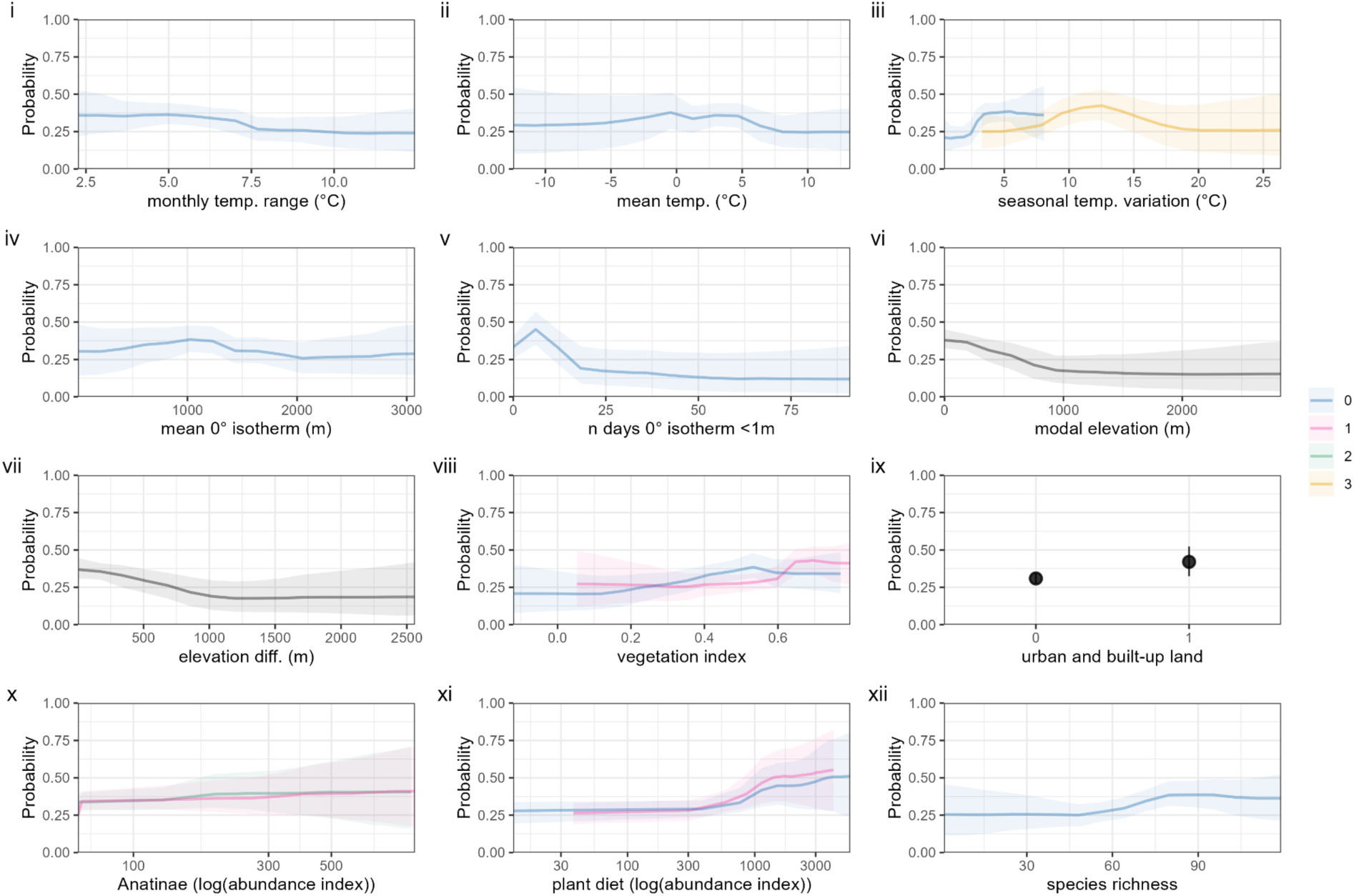
Full partial dependence of BART model trained on dataset B, non-breeding season. Partial dependence associated with all covariates in final BART model of aggregated non-breeding season (30th November - 28th February) fitted to geospatialHPAI data for period B (10/8/2021 to 29/2/2024, ongoing H5N1 outbreak). Y axis denotes marginal probability of HPAI presence, i.e., averaging out effects of all other covariates. For continuous covariates, solid lines denote median values while shaded areas denote uncertainty via 2.5th percentile and 97.5th percentile values over 8000 draws from the posterior tree space. For categorical covariates, points denote median values while error bars denote via 2.5th percentile and 97.5th percentile values. Colours denote seasonal delay increasing from 0 (predicted season) to 3 (three seasons prior to predicted season); black denotes non-seasonally-variable covariates.

**Supplemental Figure S7.**
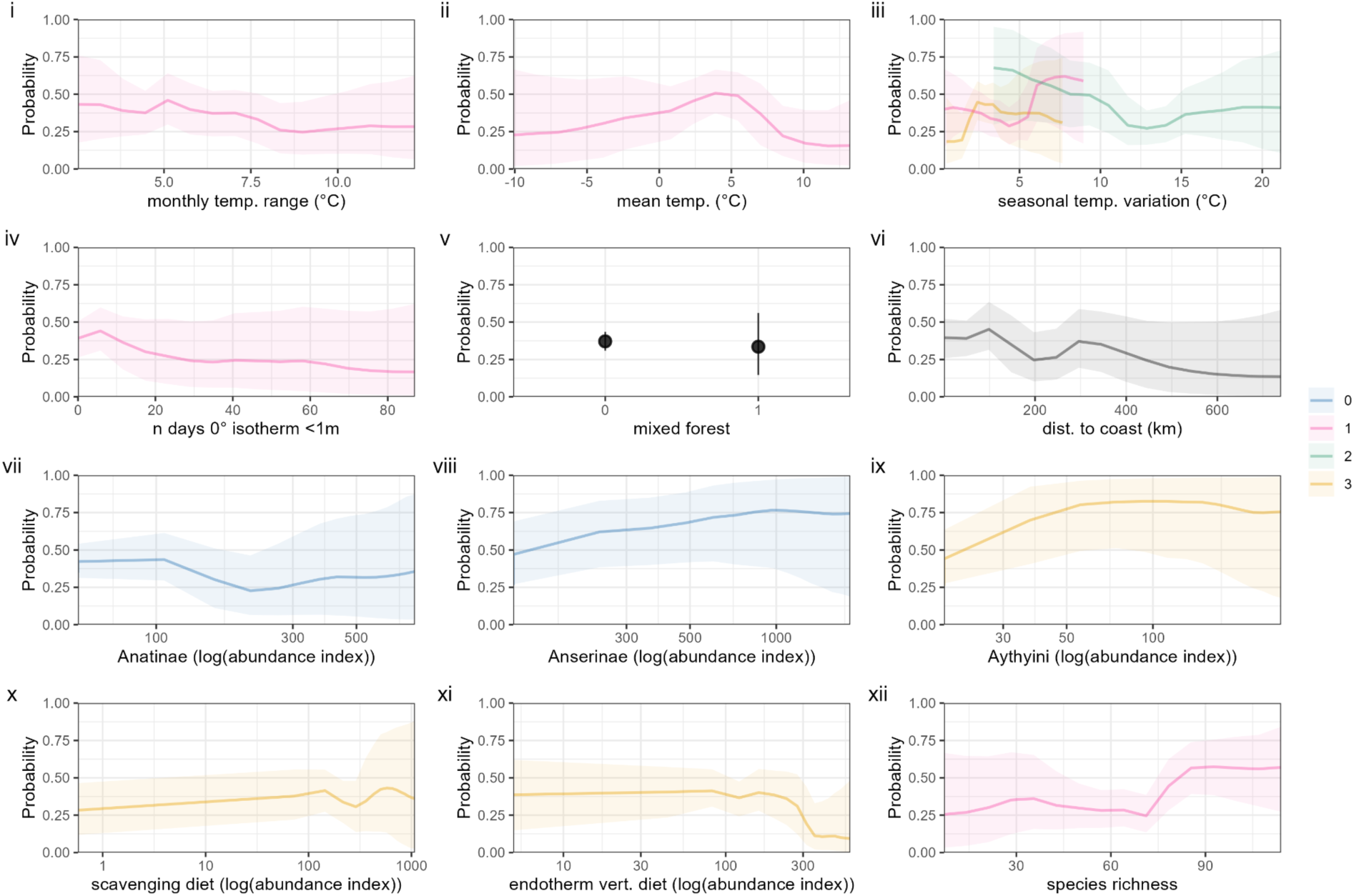
Full partial dependence of BART model trained on dataset B, pre-breeding migration. Partial dependence associated with all covariates in final BART model of aggregated pre-breeding migration season (1st March - 6th June) fitted to geospatial HPAI data for period B (10/8/2021 to 29/2/2024, ongoing H5N1 outbreak). Y axis denotes marginal probability of HPAI presence, i.e., averaging out effects of all other covariates. For continuous covariates, solid lines denote median values while shaded areas denote uncertainty via 2.5th percentile and 97.5th percentile values over 8000 draws from the posterior tree space. For categorical covariates, points denote median values while error bars denote via 2.5th percentile and 97.5th percentile values. Colours denote seasonal delay increasing from 0 (predicted season) to 3 (three seasons prior to predicted season); black denotes non-seasonally-variable covariates.

**Supplemental Figure S8.**
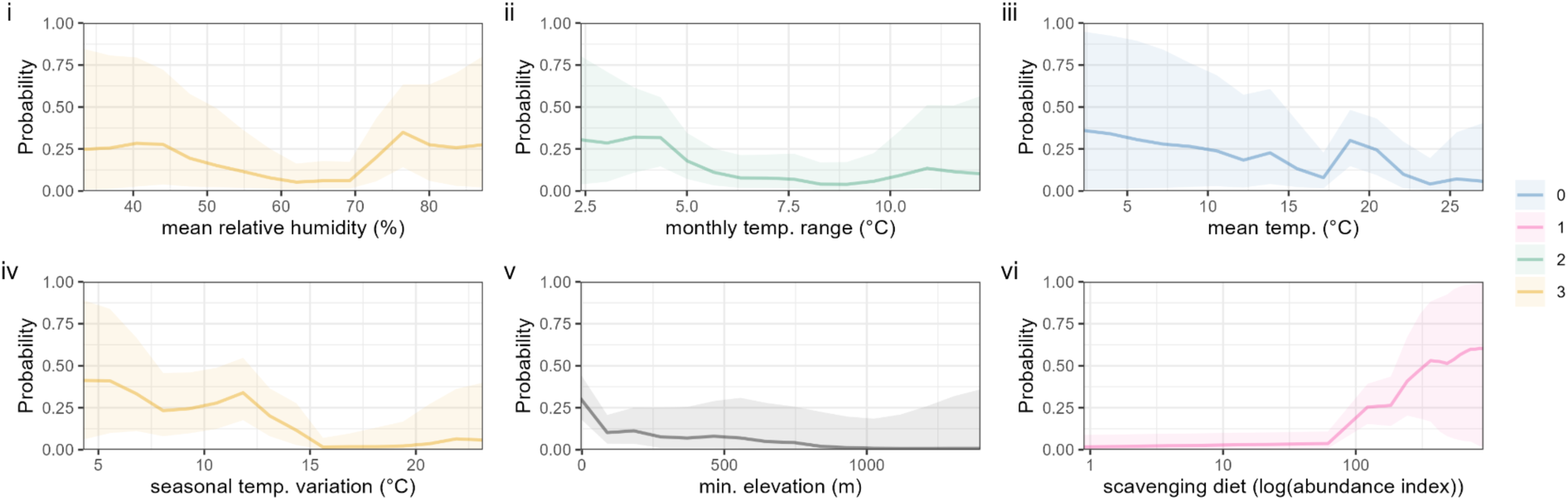
Full partial dependence of BART model trained on dataset B, breeding season. Partial dependence associated with all covariates in final BART model of aggregated breeding season (7th June - 9th August) fitted to geospatial HPAI data for period B (10/8/2021 to 29/2/2024, ongoing H5N1 outbreak). Y axis denotes marginal probability of HPAI presence, i.e., averaging out effects of all other covariates. For continuous covariates, solid lines denote median values while shaded areas denote uncertainty via 2.5th percentile and 97.5th percentile values over 8000 draws from the posterior tree space. For categorical covariates, points denote median values while error bars denote via 2.5th percentile and 97.5th percentile values. Colours denote seasonal delay increasing from 0 (predicted season) to 3 (three seasons prior to predicted season); black denotes non-seasonally-variable covariates.

**Supplemental Figure S9.**
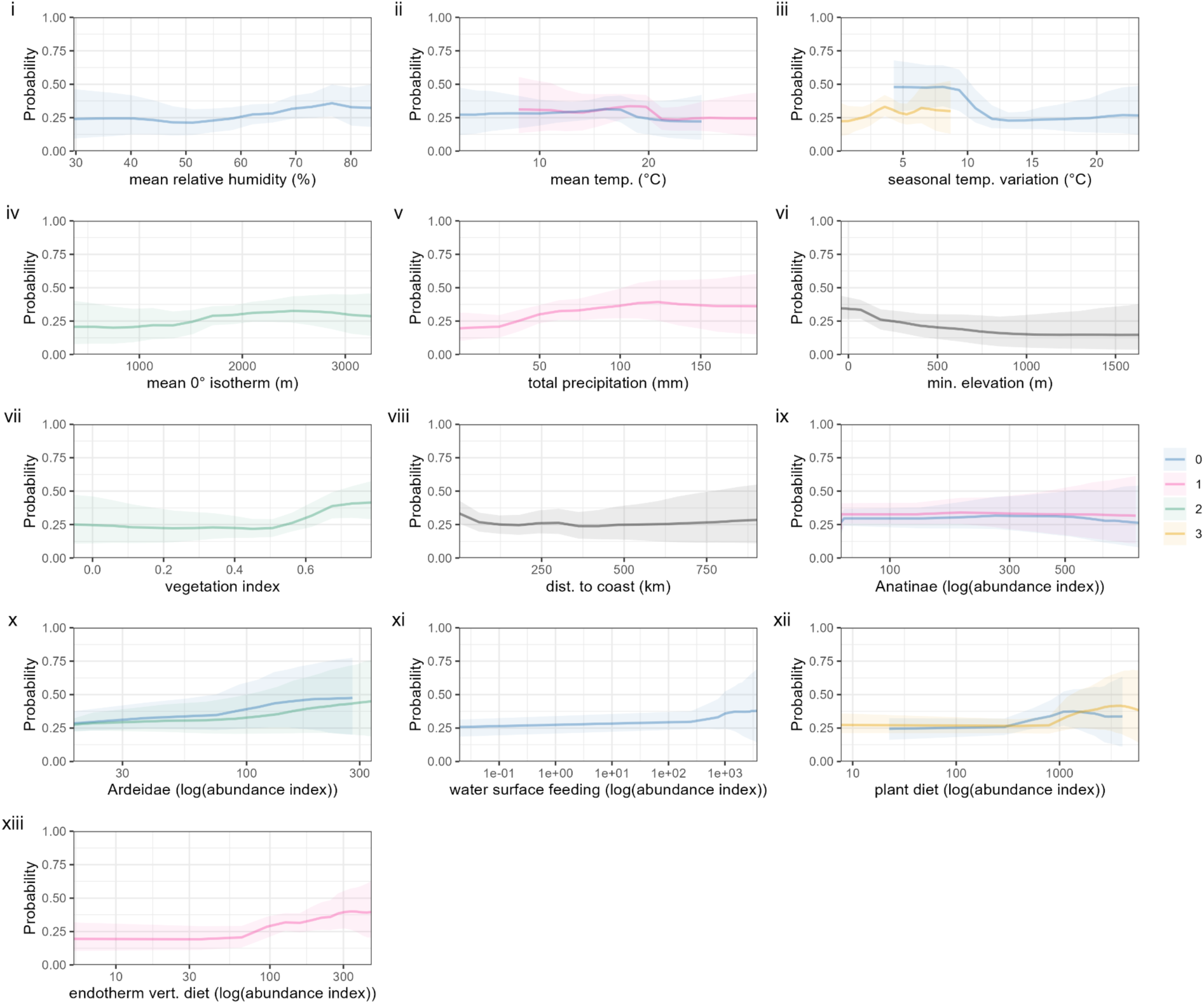
Full partial dependence of BART model trained on dataset B, post-breeding migration. Partial dependence associated with all covariates in final BART model of aggregated post-breeding migration season (10th August - 29th November) fitted to geospatial HPAI data for period B (10/8/2021 to 29/2/2024, ongoing H5N1 outbreak). Y axis denotes marginal probability of HPAI presence, i.e., averaging out effects of all other covariates. For continuous covariates, solid lines denote median values while shaded areas denote uncertainty via 2.5th percentile and 97.5th percentile values over 8000 draws from the posterior tree space. For categorical covariates, points denote median values while error bars denote via 2.5th percentile and 97.5th percentile values. Colours denote seasonal delay increasing from 0 (predicted season) to 3 (three seasons prior to predicted season); black denotes non-seasonally-variable covariates.

**Supplemental Figure S10.**
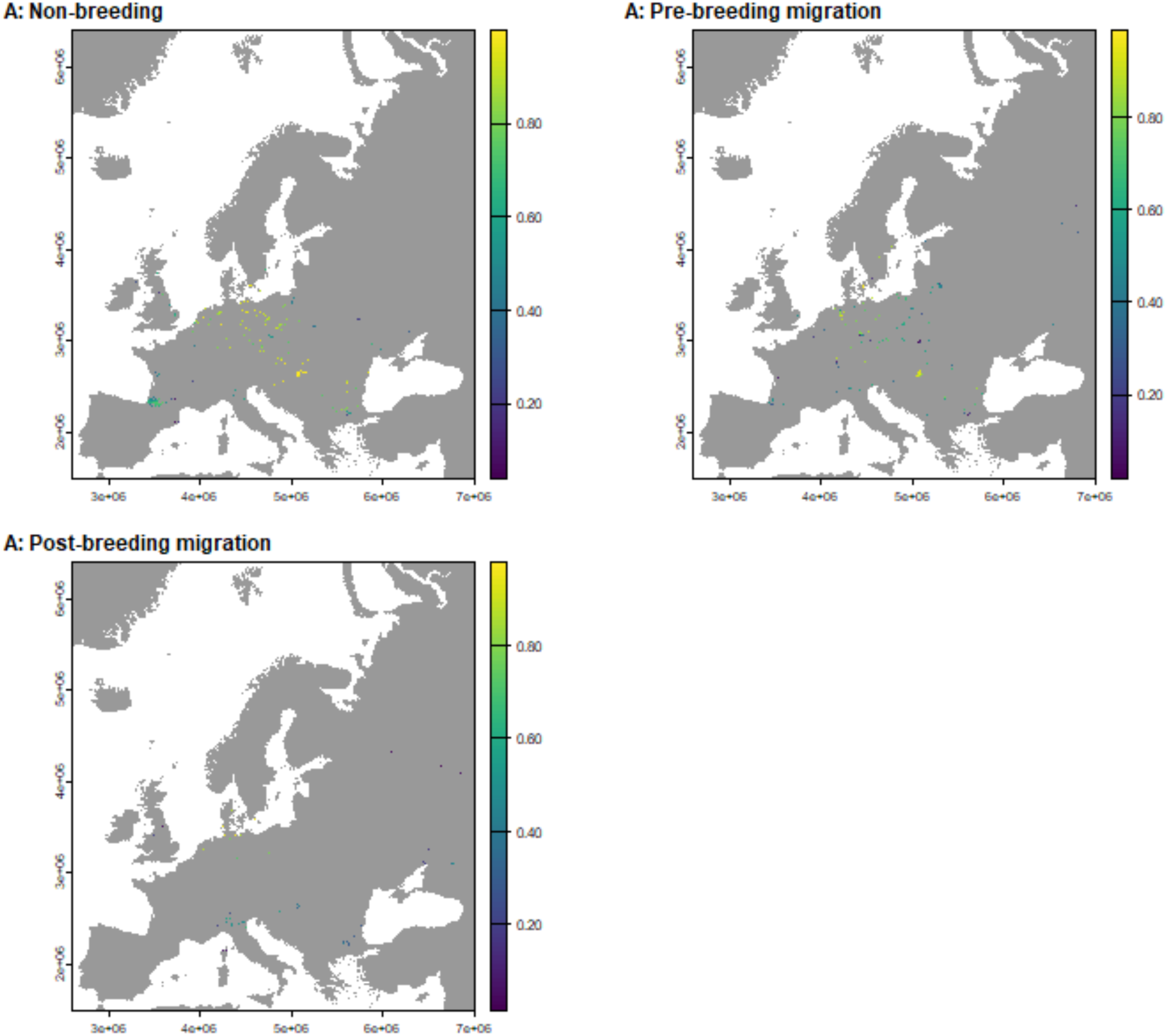
Domestic bird cases and model-predicted wild bird risk, period A. Maps of the study area with cells containing HPAI in domestic birds highlighted and coloured by the predicted probability from BART models over period A (10/8/2016 to 9/8/2021, spanning H5N8 and H5N6 events). Panels denote individual seasons.

**Supplemental Figure S11.**
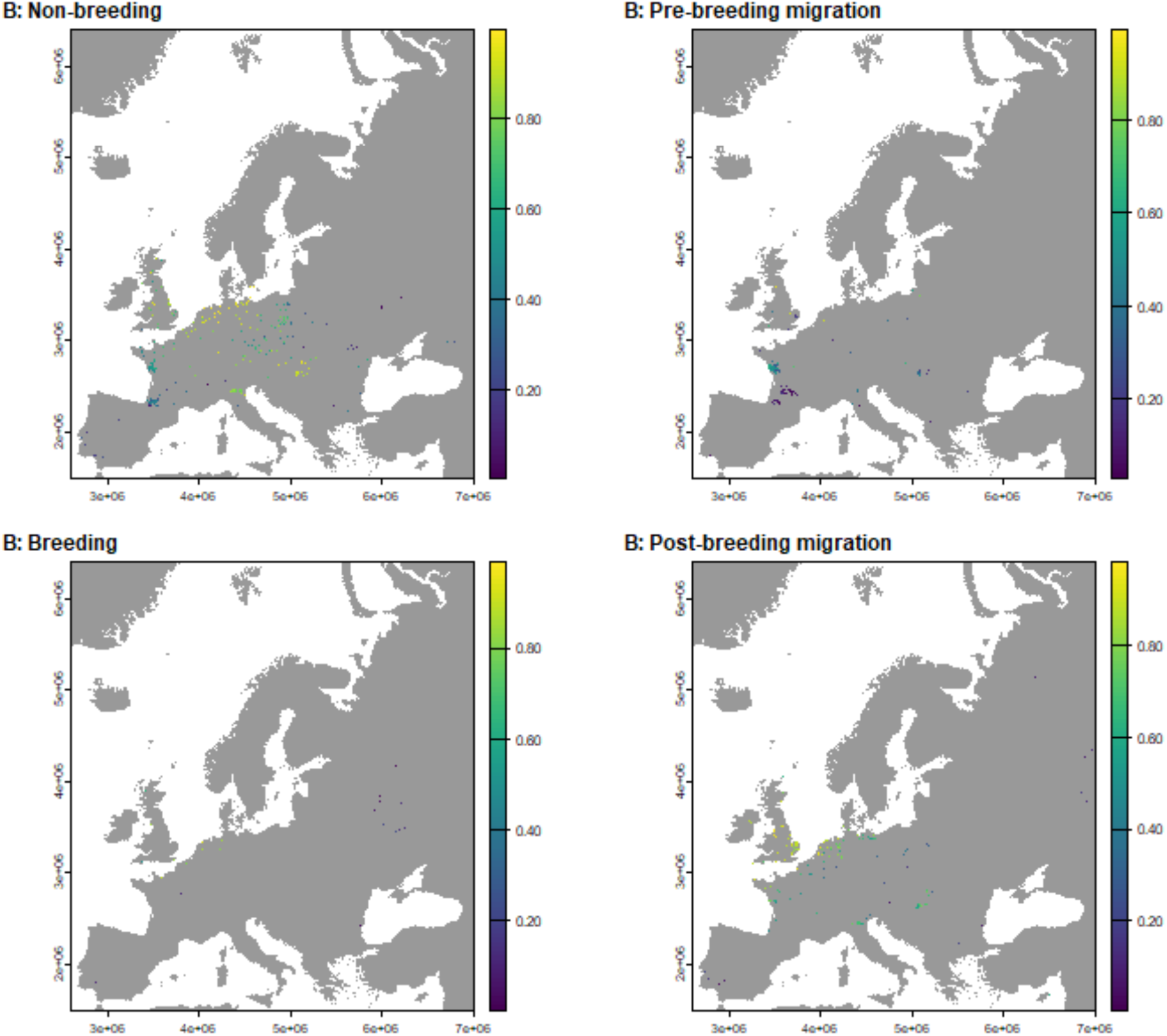
Domestic bird cases and model-predicted wild bird risk, period B. Maps of the study area with cells containing HPAI in domestic birds highlighted and coloured by the predicted probability from BART models over period B (10/8/2021 to 29/2/2024, ongoing H5N1 outbreak). Panels denote individual seasons.

## Supplemental Methods S1

### Rationalising taxonomic naming

Taxonomic labelling was not consistent between our species-level data sources, meaning that certain species identified by eBird as being present in Europe did not appear in some of the other data sources, despite these sources theoretically being more comprehensive than the eBird Status and Trends data (which is limited to those species which are confidently modelled by eBird). To remedy this we performed a semi-automated rationalisation process using the AVONET database of bird species names^72^. Each datapoint in the AVONET database corresponds to a single species and includes binomial names from three different resources (BirdLife, eBird, and BirdTree) as well as an alphanumeric ID string, allowing for identification of synonyms when comparing data from sources using differing taxonomies. For each species identified by eBird as being present in Europe (obtained by filtering by region within the eBird status and trends data) we used AVONET to generate a set of possible synonyms for that species. We matched eBird species abundance records of a species with species-level trait data from other datasets by first searching for that species’ binomial name in that dataset, and then searching for its synonyms if the binomial name was missing. Where the synonyms were absent or where multiple synonyms for a species appeared in the trait data, we checked the data manually and used a combination of renaming and removal of problematic species to ensure that each species had exactly one match in each trait dataset, based on the Avibase online bird taxonomy database ^73^.

Our species-level trait data construction begins by loading in 708 species records from eBird Status and Trends. We then bring in the list of global species population size estimates from Callaghan *et al.* ^74^, and for each species in the eBird Status and Trends data attempt to assign a population size. The following 6 species in the eBird list could not be matched to any records in the population size estimate data:

● Northern Hawk Owl, *Surnia ulula*
● Iberian Green Woodpecker, *Picus sharpei*
● Western Subalpine Warbler, *Curruca iberiae*
● Kruper’s Nuthatch, *Sitta krueperi*
● Amur Stonechat, *Saxicola stejnegeri*
● Eastern Black-eared Wheatear, *Oenanthe melanoleuca*

We removed *Surnia ulula*, *Sitta krueperi*, and *Saxicola stejnegeri* from our species list as we were unable to find possible matches in the population size estimate data. *Picus sharpei* and *Oenanthe melanoleuca* are classified in some sources as subspecies of other species found in eBird (*Picus viridis*, the European green woodpecker, and *Oenanthe hispanica*, the Western black-eared wheatear, respectively). In these cases it was necessary to remove both species in each pair from our species list because the total *Picus viridis* population size in Callaghan *et al.* effectively included both *Picus viridis* and *Picus hispanica*, meaning that keeping *Picus viridis* in the model would result in us effectively overestimating the numbers of *Picus viridis* present outside of *Picus hispanica*’s range. The same considerations motivated us to remove both *Oenanthe melanoleuca* and *Oenanthe hispanica*. The genus *Curruca* has only recently been recognised and does not appear in some sources and so we remove all of its members. After these removals we were left with 691 of the initial 708 species. Checking the CLOVER database with scientific names and pseudonyms from AVIBASE revealed that none of the species we removed were known avian influenza hosts.

After matching the species from eBird Status and Trends to population size estimates, we used the AVONET database to identify possible species synonyms. The following species could not be found in AVONET:

● Grey-headed Swamphen, *Porphyrio poliocephalus*
● Yellow-headed caracara, *Daptrius chimachima*
● Oriental cuckoo, *Cuculus optatus*

Consulting eBird suggested that *Porphyrio poliocephalus* is found only in small numbers on the outskirts of Europe ^99^, and so we removed it from our species list. We replaced *Daptrius chimachima* with *Milvago chimachima*, which is synonymous with *Daptrius chimachima* ^100^. Although *Cuculus optatus* was absent from AVONET, it was present in all of our other databases meaning we were able to match it to species-level traits without using synonyms, and so we kept it in our species list without renaming. This left us with 690 of our initial 708 species.

The binomial names for three of the known host species in CLOVER could not be found in AVONET, motivating the following changes to binomial names in CLOVER based on species entries in Avibase ^73^:

● Eurasian jackdaw, *Coloeus monedula*, replaced with *Corvus monedula;*
● Mongolian gull, *Larus mongolicus*, replaced with *Larus cachinnans;*
● Little cuckoo, *Piaya minuta*, replaced with *Coccycua minuta*.

We replaced the binomial names of the following four species in EltonTraits to ensure that every species from EltonTraits could be matched with a single species from eBird Status and Trends:

● Long-tailed pipit, *Anthus longicaudatus*, replaced with *Anthus vaalensis*
● Iquitos gnatcatcher, *Polioptila clementsi*, replaced with *Polioptila guianensis*
● Bluntschli’s vanga, *Hypositta perdita*, replaced with *Oxylabes madagascariensis*
● Vietnamese pheasant, *Lophura hatinhensis*, replaced with *Lophura edwardsi*
● Yellow warbler, *Setophaga petechia*, replaced with *Dendroica petechia*

In the IUCN trait data we changed the binomial name of the yellow-throated parrotbill from *Suthora webbiana* to *Sinosuthora webbiana*.

### Derivation of behavioural season boundaries

The species metadata available through eBird Status and Trends includes fields listing the start and end dates of the breeding, nonbreeding, post-breeding migration, and pre-breeding migration seasons for species which exhibit these seasons ^59^. These are listed as calendar dates during the calendar year 2021, when the data underlying the 2022 release of the Status and Trends data was collected. These dates are to a resolution of one week, so that each date occurred on a Monday and “consecutive” dates are 7 days apart. Not all species have season start and end dates listed for all four behavioural seasons, with some species having no dates listed or dates listed for a subset of the four seasons. However, we were able to directly verify that for each species with start and end dates present for at least one season, the times between the listed dates cover the entire calendar year; equivalently, the list of species with a behavioural season listed for any given day is consistent throughout the year.

Of the 708species listed as being present in Europe in eBird Status and Trends, 546 species had behavioural season start and end dates listed. For each of these 546 species, we used the season start and end dates listed in eBird Status and Trends to assign a behavioural season for each day of the calendar year 2021. We then calculated the total number of species in each behavioural season for each day of the year to give a daily frequency distribution of species in each behavioural season. This time-varying distribution is plotted in Supplemental Figure S1A. Visual inspection shows that at the cross-species level the dominant behavioural-seasonal trends move in waves, with successive periods where the majority of species are in each of the successive behavioural seasons. To define cross-species season start and end dates, we calculated the first date for each behavioural season on which a plurality of species are in that behavioural season; that is, the first date on which the number of species in that behavioural season is larger than the number in any other season (note that since the majority of species are in the nonbreeding season at the start of calendar year 2021, we defined the start of the nonbreeding season to be the day after the last day on which a plurality of species were in the post-breeding season; visual inspection of the daily frequency distributions confirms that these two seasons do indeed follow each other at the cross-species level). The resulting behavioural season start/end dates were: non-breeding season, 30th November - 28th February (calendar days 334 - 59); pre-breeding migration, 1st March - 6th June (days 60 - 157); breeding season, 7th June - 9th August (days 158 - 221); and post-breeding migration, 10th August - 29th November (days 222 - 333). For data from leap years in our model training and testing, we considered 29th February as part of the non-breeding season.

### Random intercept models

We attempted to augment both the basic and cross-seasonal models by specifying a random intercept according to country. We labelled each raster cell by the country its coordinates lie within using the R package rworldmap, v1.3.8 ^101^, and used the random intercept option provided in *embarcadero* to fit the random intercept term. For each model we ran a single MCMC chain to generate 1000 posterior samples. The random intercept term is intended to capture country-level heterogeneities in HPAI detection and reporting capacity. However, since our pseudo-absence generation process accounts for spatial heterogeneity in observation processes through its weighting on citizen bird observation intensity data, there is a possibility that introducing a random intercept term may be “double accounting” for spatial heterogeneity, motivating us to compare models with and without random intercept. Along with this double accounting problem, the substantial spatial heterogeneity in our data means that the testing data for the different dataset-season combinations consistently contains data from countries which do not appear in the corresponding training data, which is likely to make assessment of model performance difficult.

Performance metrics for the models with random intercept included are listed in Supplemental Table S1. Comparison with Table 4 of the main text suggests that the random intercept models do not offer a substantial improvement in predictive ability over the corresponding models without random intercept terms, sensitivity being notably weaker for period B models.

**Supplemental Table S1.**
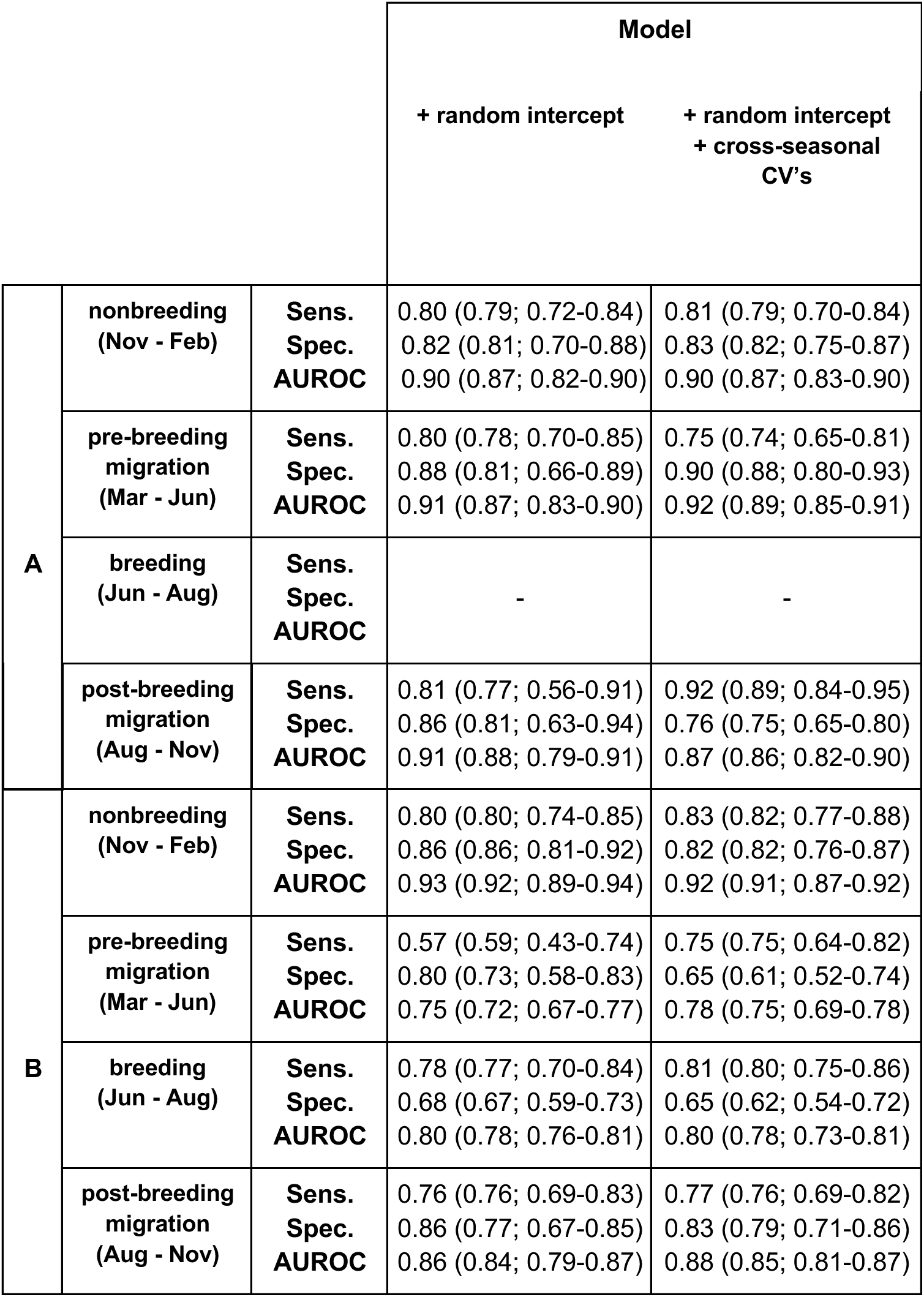
Posterior means of performance metrics for random intercept models on each dataset’s test set. AUROC denotes Area Under the Receiver-Operating Characteristic curve, CV denotes covariates. Presence data in test set A is from the 2020-2021 H5N8 HPAI clade 2.3.4.4b outbreak. Presence data in test set B is from 1/3/2023 to 29/2/2024, during the ongoing (since 2021) H5N1 HPAI clade 2.3.4.4b outbreak. Values in brackets denote posterior median and 95% credible interval of metrics within individual trees. As posterior mean values are based on consensus predictions averaging over all sampled trees, this can exceed posterior medians and 95% credible intervals.

|  |  |  | **Model** |  |
| --- | --- | --- | --- | --- |
|  |  |  | **+ random intercept** | **+ random intercept**<br>**+ cross-seasonal**<br>**CV's** |
| **A** | **nonbreeding**<br>**(Nov - Feb)** | **Sens.**<br>**Spec.**<br>**AUROC** | 0.80 (0.79; 0.72-0.84)<br>0.82 (0.81; 0.70-0.88)<br>0.90 (0.87; 0.82-0.90) | 0.81 (0.79; 0.70-0.84)<br>0.83 (0.82; 0.75-0.87)<br>0.90 (0.87; 0.83-0.90) |
|  | **pre-breeding**<br>**migration**<br>**(Mar - Jun)** | **Sens.**<br>**Spec.**<br>**AUROC** | 0.80 (0.78; 0.70-0.85)<br>0.88 (0.81; 0.66-0.89)<br>0.91 (0.87; 0.83-0.90) | 0.75 (0.74; 0.65-0.81)<br>0.90 (0.88; 0.80-0.93)<br>0.92 (0.89; 0.85-0.91) |
|  | **breeding**<br>**(Jun - Aug)** | **Sens.**<br>**Spec.**<br>**AUROC** | - | - |
|  | **post-breeding**<br>**migration**<br>**(Aug - Nov)** | **Sens.**<br>**Spec.**<br>**AUROC** | 0.81 (0.77; 0.56-0.91)<br>0.86 (0.81; 0.63-0.94)<br>0.91 (0.88; 0.79-0.91) | 0.92 (0.89; 0.84-0.95)<br>0.76 (0.75; 0.65-0.80)<br>0.87 (0.86; 0.82-0.90) |
| **B** | **nonbreeding**<br>**(Nov - Feb)** | **Sens.**<br>**Spec.**<br>**AUROC** | 0.80 (0.80; 0.74-0.85)<br>0.86 (0.86; 0.81-0.92)<br>0.93 (0.92; 0.89-0.94) | 0.83 (0.82; 0.77-0.88)<br>0.82 (0.82; 0.76-0.87)<br>0.92 (0.91; 0.87-0.92) |
|  | **pre-breeding**<br>**migration**<br>**(Mar - Jun)** | **Sens.**<br>**Spec.**<br>**AUROC** | 0.57 (0.59; 0.43-0.74)<br>0.80 (0.73; 0.58-0.83)<br>0.75 (0.72; 0.67-0.77) | 0.75 (0.75; 0.64-0.82)<br>0.65 (0.61; 0.52-0.74)<br>0.78 (0.75; 0.69-0.78) |
|  | **breeding**<br>**(Jun - Aug)** | **Sens.**<br>**Spec.**<br>**AUROC** | 0.78 (0.77; 0.70-0.84)<br>0.68 (0.67; 0.59-0.73)<br>0.80 (0.78; 0.76-0.81) | 0.81 (0.80; 0.75-0.86)<br>0.65 (0.62; 0.54-0.72)<br>0.80 (0.78; 0.73-0.81) |
|  | **post-breeding**<br>**migration**<br>**(Aug - Nov)** | **Sens.**<br>**Spec.**<br>**AUROC** | 0.76 (0.76; 0.69-0.83)<br>0.86 (0.77; 0.67-0.85)<br>0.86 (0.84; 0.79-0.87) | 0.77 (0.76; 0.69-0.82)<br>0.83 (0.79; 0.71-0.86)<br>0.88 (0.85; 0.81-0.87) |

